# Modular slowing of resting-state dynamic Functional Connectivity as a marker of cognitive dysfunction induced by sleep deprivation

**DOI:** 10.1101/2020.01.17.910810

**Authors:** Diego Lombardo, Catherine Cassé-Perrot, Jean-Philippe Ranjeva, Arnaud Le Troter, Maxime Guye, Jonathan Wirsich, Pierre Payoux, David Bartrés-Faz, Régis Bordet, Jill C Richardson, Olivier Felician, Viktor Jirsa, Olivier Blin, Mira Didic, Demian Battaglia

## Abstract

Dynamic Functional Connectivity (dFC) in the resting state (rs) is considered as a correlate of cognitive processing. Describing dFC as a flow across morphing connectivity configurations, our notion of dFC speed quantifies the rate at which FC networks evolve in time. Here we probe the hypothesis that variations of rs dFC speed and cognitive performance are selectively interrelated within specific functional subnetworks.

In particular, we focus on Sleep Deprivation (SD) as a reversible model of cognitive dysfunction. We found that whole-brain level (global) dFC speed significantly slows down after 24h of SD. However, the reduction in global dFC speed does not correlate with variations of cognitive performance in individual tasks, which are subtle and highly heterogeneous. On the contrary, we found strong correlations between performance variations in individual tasks –including Rapid Visual Processing (RVP, assessing sustained visual attention)– and dFC speed quantified at the level of functional subnetworks of interest. Providing a compromise between classic static FC (no time) and global dFC (no space), modular dFC speed analyses allow quantifying a different speed of dFC reconfiguration independently for sub-networks overseeing different tasks. Importantly, we found that RVP performance robustly correlates with the modular dFC speed of a characteristic frontoparietal module.

**Highlights:**

- Sleep Deprivation (SD) slows down the random walk in FC space implemented by Dynamic Functional Connectivity (dFC) at rest.
- Whole-brain level slowing of dFC speed does not selectively correlate with fine and task-specific changes in performance
- We quantify dFC speed separately for different link-based modules coordinated by distinct regional “meta-hubs”
- Modular dFC speed variations capture subtle and task-specific variations of cognitive performance induced by SD.

**Author summary:** We interpreted dynamic Functional Connectivity (dFC) as a random walk in the space of possible FC networks performed with a quantifiable “speed”.

Here, we analyze a fMRI dataset in which subjects are scanned and cognitively tested both before and after Sleep Deprivation (SD), used as a reversible model of cognitive dysfunction. While global dFC speed slows down after a sleepless night, it is not a sufficiently sensitive metric to correlate with fine and specific cognitive performance changes. To boost the capacity of dFC speed analyses to account for fine and specific cognitive decline, we introduce the notion of *modular dFC speed*. Capitalizing on an edge-centric measure of functional connectivity, which we call Meta-Connectivity, we isolate subgraphs of FC describing relatively independent random walks (*dFC modules*) and controlled by distinct “puppet masters” (*meta-hubs*). We then find that variations of the random walk speed of distinct dFC modules now selectively correlate with SD-induced variations of performance in the different tasks. This is in agreement with the fact that different subsystems – distributed but functionally distinct– oversee different tasks.

The high sensitivity of modular dFC analyses bear promise of future applications to the early detection and longitudinal characterization of pathologies such as Alzheimer’s disease.

## Introduction

The majority of studies on resting state functional brain connectivity so far have considered time averaged resting state Functional Connectivity (rs FC). However, substantial evidence suggest that the brain is restless even at rest and that rs FC continually fluctuates in a way which is far from being random, displaying non trivial spatiotemporal structures (Hutchison et al., 2013; Calhoun et al., 2014; Gonzalez-Castillo & Bandettini, 2018). Such dynamic Functional Connectivity (dFC), during both rest and task conditions, has been suggested to be an indicator of cognitive processing (Cohen, 2018). Beyond differences of age (Hutchison & Morton, 2015) and gender (Yaesoubi et al., 2015), or changes in vigilance (Wang et al., 2016; Shine et al., 2016) and consciousness (Hudetz et al., 2015; Cavanna et al., 2018), higher “fluidity” of dFC has been linked to higher mindfulness (Lin et al., 2018), levels of attention (Kucyi et al., 2017), faster learning (Bassett et al., 2011), flexibility of executive control (Braun et al., 2015) and, more generally, superior performance in several cognitive and behavioral domains (Jia et al., 2014). These findings are in line with a view of the human brain as a dynamical system, in which flexible communication and large-scale integration between multiple, possibly remote, local subnetworks underlies emergent information processing and cognitive operations (Mesulam et al. 1990, Varela et al., 2001; Bressler & Kelso, 2016; Kirst et al., 2016).

The shift to a dynamic chronnectome (Calhoun et al., 2014) perspective is however hampered by the lack of a consensus on the best approach to estimate not-artefactual dFC to be chosen amongst an increasingly larger spectrum of diverse methods (Preti et al., 2017). In particular, statistical concerns have been raised: firstly, about whether resting-state FC is really non-stationary (Zaleski et al., 2014; Hindriks et al., 2016) and, secondly, whether discrete connectivity states do exist or can be reliably extracted (Shakil et al., 2016; Liégeois et al., 2017). Here we will advance on investigating relations between dFC and cognitive performance, by capitalizing on an original methodological framework which circumvents both these concerns. In Battaglia et al. (2020) we introduced the notion of *dFC speed*, quantifying in a time-resolved manner the rate at which brain-wide FC networks are changing from one time-window to the next. We thus interpret dFC as a random walk in the highdimensional space of possible FC network realizations, describing it as a smooth flow across continually morphing connectivity configurations, rather than as a non-stationary sequence of sharp inter-state transitions. In this study, we go beyond Battaglia et al. (2020) by exploring with more precision and detail the hypothesis that variations in resting state dFC speed correlate with cognitive performance. Specifically, we substantially extend the approaches of Battaglia et al. (2020) by quantifying dFC changes not only at the whole-brain level but also at the level of specific functional subnetworks, believing that these modular changes are better at capturing fine cognitive performance variations across specific tasks.

We concentrate, as a proof of concept, on *Sleep Deprivation (SD)* as a model of cognitive dysfunction, useful to understand how changes of dFC correlate with reversible cognitive impairment. Furthermore, SD is a common condition in modern societies with a significant impact on productivity and health (Institute of Medicine Committee on Sleep and Research, 2006), affecting cognitive functions such as executive control and attention (Durmer & Dinges, 2005; Krause et al., 2017). Finally, investigating dFC in SD may lead to the development of novel predictive markers of early network dysfunction in disease. Indeed, SD has also been proposed as a cognitive challenge model of cognitive impairment arising in Alzheimer’s Disease (AD), because of both the partially overlapping spectrum of induced cognitive deficits and their response to pharmacological treatment (Cheeh & Chuah, 2008; Repantis et al., 2010; Wirsich et al., 2018). For this reason, cognitive tests chosen with a sensitivity to both SD and cognitive dysfunction in AD were chosen for the present study (Cummings et al. 2012, Randall et al. 2005, Dodds et al. 2011, Kirova et al. 2015).

Previous investigations have shown that SD affects time-averaged FC at the whole-brain level (Yeo, Tandi & Chee, 2015; Kaufmann et al., 2016; Wirsich et al., 2018). However, it is not very clear how such changes of static network topology relate to cognitive effects of sleep deprivation. Other studies considered classic dFC analyses and found that transition dynamics between global FC states are slowed down after SD and during descent from wakefulness into sleep (El-Baba et al., 2019). Furthermore changes in state transitions and occupancy could predict the rate of decline of speed of processing and working memory after multiple consecutive nights of SD (Patanaik et al., 2018). Here we go beyond these promising results by studying how 24 hs of SD affect our novel dFC speed markers in relation to cognitive performance looking at changes at not only the global but also the subnetwork level.

Firstly, we observe that the speed of reconfiguration of resting-state whole-brain FC networks (*global dFC speed*) is significantly slowed-down by SD. Secondly, we demonstrate a correlation between the modular dFC speed of a specific frontoparietal sub-network –related to well-known attentional networks (Corbetta & Shulman, 2002)– and performance on a demanding cognitive test, Rapid Visual Processing (RVP), assessing sustained visual attention and central executive control (Coull et al., 1996).

In Battaglia et al. (2020) we found that higher dFC speeds at the whole-brain level were associated with higher levels of general cognitive performance. Here, however we refine our analyses going beyond rough cognitive assessments –as the MOCA score (Nasreddine et al., 2005) used in Battaglia et al. (*submitted as a companion paper*)–, optimized for diagnosis but able just to detect sizeable cognitive deficits affecting several cognitive domains simultaneously. The effects of SD can on the contrary be very moderate and diverse for different tasks, requiring a much finer sensitivity and selectivity of analysis. Here, we find then that reductions of global dFC speed at the whole brain level do not predict these small and specific changes of cognitive performance in different individual tasks. Importantly, however, we observe again significant correlations with detailed changes in cognitive performance after introducing a novel type of metric, which we call *modular dFC speed*. Unlike global dFC speed analyses – in which information about *where* network changes are happening, because the rate of change is averaged over the whole brain are lost – modular dFC speed allows for the quantification of a different speed of dFC reconfiguration independently for each different subnetwork of interest. Correlations with cognitive performance are then highlighted when focusing on task-relevant networks, which we extract in an unsupervised manner –through an analysis of *Meta-Connectivity* (MC), i.e. correlation between time-dependent functional links (cf. Bassett et al., 2014, Brovelli et al., 2017; Faskowitz et al., 2019)–and whose independent dFC fluctuations would have otherwise been undetectable by whole-brain dFC speed analyses.

Our findings suggest that analyses of dFC speed variations are well able to detect the temporal reorganization of resting state FC fluctuations after SD. Furthermore, beyond mere changes in reaction time or vigilance, our quantifications of dFC variations robustly track the magnitude of changes in cognitive performance induced by SD within each specific subject. Importantly, modular dFC speed measures are able to detect subtle changes in dFC that are confined to specific sub-networks. This makes it possible to capture variations of performance in cognitive function such as attention – relying on the interaction between spatially well-defined systems (Corbetta & Shulman, 2002).

## Material and methods

### Participants

We here re-analyze neuropsychological and fMRI data previously reported by Wirsich et al. (2018). Within the framework of a work-package of the EU PharmaCog Innovative Medicine Initiative (https://www.imi.europa.eu/projects-results/project-factsheets/pharma-cog), fifteen healthy subjects were recruited (all male, 25–40 years old, Body Mass Index, BMI ≤ 27, right-handed) to participate in an experiment distributed over ten different days in blocks of two days each. Before inclusion, volunteers were checked for their medical history, physical examination, vital signs, results of blood chemistry and haematology, urine drug screen, electrocardiogram (ECG) and their visual and auditory abilities. Volunteers who received medical treatment, smoked more than five cigarettes per day, took more than five caffeine or energy drinks per day or suffered from claustrophobia were not included. It was ensured that the volunteers had good regular sleep habits and rhythms using questionnaires and clinical interviews (Pittsburgh Quality Index with regular sleep between 6.5 and 9 h per night, Epworth scale score below 10, Horne and Ostberg scale score above 31 and below 69, absence of jet lag and no occupation with time-shifts), no history of clinicalsigns of sleep disorder (Berlin sleep apnea scale, Restless Legs Syndrome questionnaire) and no history of psychiatric disorder (psychiatric interview with the M.I.N.I-DSM IV). Three days before the session, the subjects wore an actigraph and completed a sleep diary to verify their sleep status. Four subjects quit the protocol before the end (three withdrawals of consent and one drop-out for an adverse event), which resulted in a final group of eleven subjects (age=34±4 years, range 28–40 years).

In the original experiment, some of the subjects were administered doses of various neuroenhancer drugs prior to the fMRI scan and cognitive testing. For more details, see Wirsich et al (2018) and Chan Kwong et al. (2020). Here, however, we prefer to focus uniquely on the placebo sessions, being more interested in statistical comparisons between dFC speed in different modules, than comparisons between dFC speed in different medication conditions. The small number of subjects does not provide sufficient statistical power to study all possible comparisons (modules *and* drugs). Therefore, for homogeneity, we consider data acquired during the placebo sessions within the first block of two days. Subjects were assessed twice, once before and once after SD with one session of rs fMRI and cognitive tests. An additional subject was discarded a posteriori because he slept for most of the time during the second day fMRI session.

### Cognitive assessment

Every subject performed seven cognitive tests assessing verbal and visual episodic memory, attention, language and working memory. In this study, we focused on three computerized tests, known to be affected by SD and selected for the complementarity of the functions they probe. Hence, psychomotor speed and attention were assessed via the Simple Reaction Time (RTI) and the Rapid Visual Processing (RVP) tests from the Cambridge Neuropsychological Test Automated Battery (CANTAB^®^). For the RTI, during 10 mn, participants, while holding a button on a box, have to react as soon as possible to yellow dots that appear on the screen by releasing the button and touching the dot on the screen. In the RVP task (12 mn), digits are shown in the centre of the screen in a pseudorandom order, at the rate of 100 digits per minute. Participants are requested to detect target sequences of 3 digits by pressing a button as quickly as possible. The participant must watch for three-target sequences at the same time. Working memory was assessed through the n-back (0-back, 1-back, 2-back, 3-back…) task (Braver et al, 1997). The subject is presented with 3 blocks of 4 continuous sequences of 16 letters. The task consists of indicating when the current letter matches the one from n steps earlier following the different instructions of each sequence. We here use as performance metric reaction times and accuracies from a 3-back task (n = 3).

Differences of cognitive performance before and after SD were compared: at the group level, using Mann-Whitney-U test or Kruskal-Wallis testing of median differences; at the within-subject level, using paired Wilcoxon signed-rank test or Kruskal-Wallis testing of median being different from zero. Bonferroni correction was applied on the number of tested cognitive performance scores (ν = 5).

### fMRI data recording and processing

For a detailed description on fMRI data acquisition, see Wirsich et al (2018). All images were acquired on a Siemens Magnetom Verio 3T MRI-Scanner (Siemens, Erlangen, Germany) with structural T_1_-weighted images acquired using a MPRAGE-sequence (TR = 2300 ms, TE = 2.98 ms, 1.0 × 1.0 × 1.0 mm, 176 slices), and 200 BOLD-sensitized EPI T2*-weighted MRI images acquired using a TR of 2.7 s (3.0 × 3.0 × 3.0 mm, TE = 30 ms, 40 slices, acquisition time of 9 min 5 s). To process fMRI data we used the SPM8 toolbox (revision 4667, http://www.fil.ion.ucl.ac.uk.gate2.inist.fr/spm/software/spm8/) to slice time and spatially realign the volumes. The AAL atlas template (*N* = 86 regions, see Figure 4B for the list of enclosed regions (left and right hemisphere; Tzourio-Mazoyer et al. (2002)) was linearly transformed into the T_1_-image of each subject (FSL 5.0 FLIRT, http://fsl.fmrib.ox.ac.uk/fsl/fslwiki/). The aligned T1-image and AAL template were then co-registered with the T2*-images (SPM8). Average cerebrospinal fluid (CSF) and white matter signal were extracted out of the T2*-images from a manually defined spherical ROI (Marsbar Toolbox 0.43, http://marsbar.sourceforge.net/). Movement parameters, CSF and white matter signals were regressed out of the fMRI time-courses extracted from each regional volume (average of all voxels). The time-series were then band-passed filtered in the 0.02 - 0.2 Hz range. Simultaneous EEG was acquired during fMRI scan and used to perform segmentation into fMRI epochs belonging to different sleep states. The resulting regional BOLD timeseries were used as input to further FC and dFC analyses. Simultaneous EEG 64 electrode EEG-cap (BrainCap-MR 3-0, Easycap, Hersching, Germany, using 63 channels in the 10–20 montage plus one extra ECG channel, reference placed on the mid-frontal position FCz) was acquired during fMRI sessions as described by Wirsich et al. (2018). EEG data was manually sleep staged for consecutive 30 s blocks by an expert (Awake, N1, N2, N3, according to the AASM sleep scoring scheme, http://www.aasmnet.org/). The acquired data was then manually sleep staged for consecutive 30 s blocks by an expert (Awake, N1, N2, N3, according to the AASM sleep scoring scheme, http://www.aasmnet.org/).

**Figure 1.**
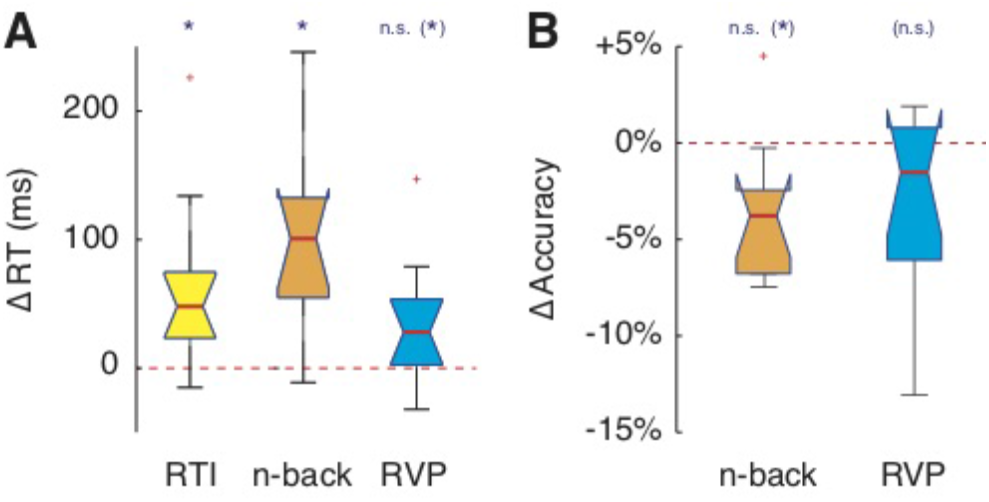
Sleep Deprivation affects cognitive performance. We tested cognitive performance in a variety of different tests including simple Reaction Time (RTI), working memory (n-back) and Rapid Visual Processing (RVP). We then quantified variations of performance (reaction times and accuracy) between before and after 24h of Sleep Deprivation (SD). **A.** Within-subject variations of RTI, n-back and RVP reaction times, significant only for RTI and n-back. For all three tests, reaction times tended to increase. **B**. Within-subject variations of n-back and RVP accuracies. Accuracy generally decreases, however only tendentially. Stars denote significancy of effect (under Wilcoxon signed rank paired test): n.s., not significant; *, *p* < 0.05; Bonferroni-corrected values outside brackets and Bonferroni-uncorrected within brackets (tendential variations).

**Figure 2.**
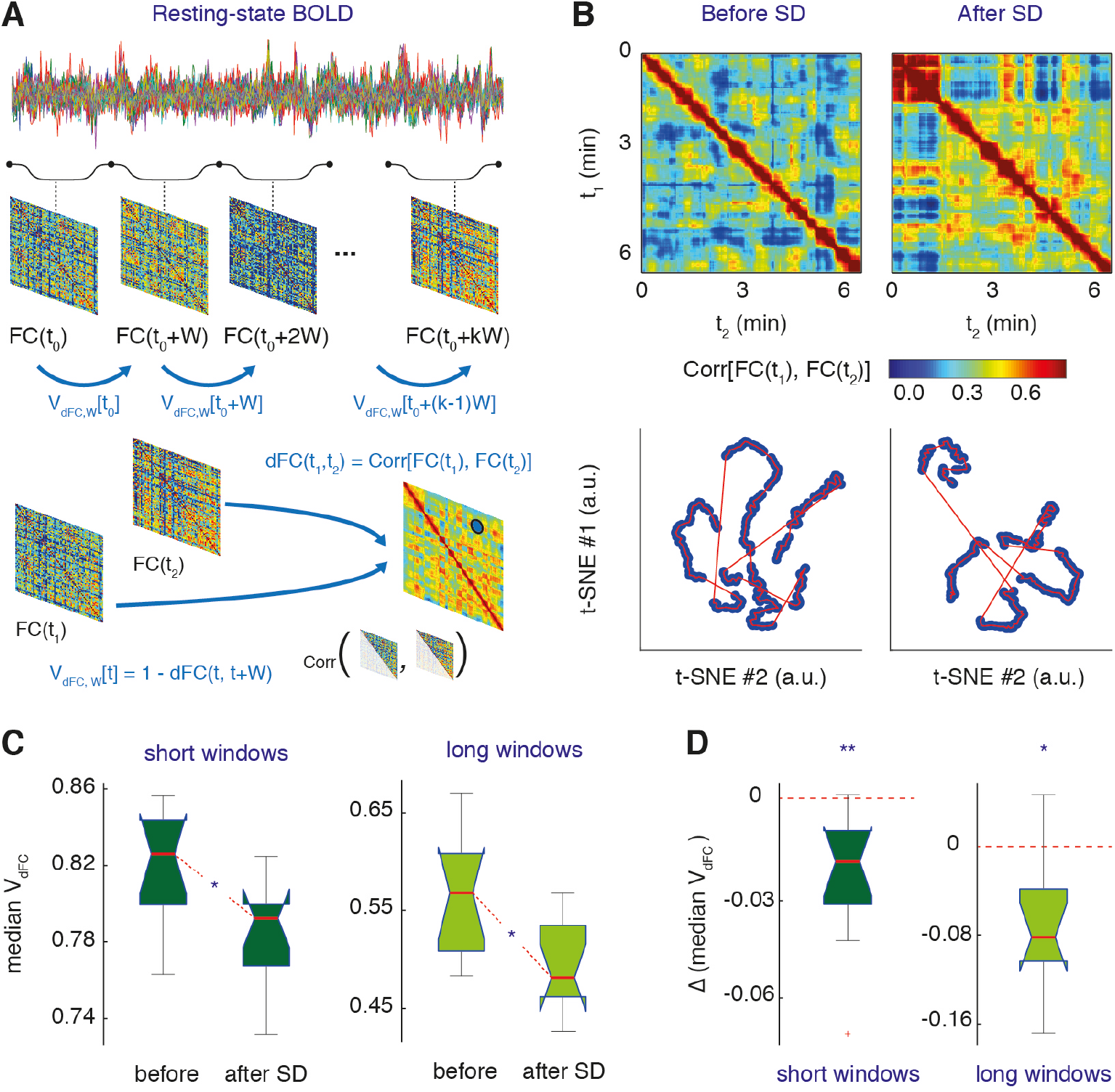
Sleep Deprivation slows down global dynamic Functional Connectivity (dFC). **A.** We reveal changes of resting state Functional Connectivity (FC) by adopting a sliding window estimation of time-resolved BOLD correlations FC(*t*). We compile the similarity between whole-brain FC matrices observed at different times into a *dFC matrix*, whose (*t_1_, t_2_*) provides the correlation between FC(*t_1_*) and FC(*t_2_*). Analogously, we evaluate *global dFC speeds V_dFC_*, i.e. the rate of reconfiguration of wholebrain FC matrices, by measuring the decorrelation between FC(*t*) matrices separated by a fixed timeinterval (equal to the chose window size *W*). **B**. On top, dFC matrices for a resting state fMRI session before SD and a second resting state fMRI session after SD for a representative subject (window size of ~40 s). A stochastic alternation between “knots” of transient FC stabilization and “leaps” of fast FC reconfiguration is visible in both cases. Below, each FC(*t*) matrix is projected into a bi-dimensional space by using a non-linear distance preserving t-SNE projection, in such a way to visualize the “random walk” followed by FC(*t*) while it reconfigures along time corresponding to the dFC matrices above. **C-D**. The median global dFC speed *V_dFC_* decreases after 24h of SD, as visible both from intergroup comparisons (panel **C**) and within-subject variations (panel **D**), for both long and short window sizes. Stars denote significancy of effect: *, *p* < 0.05; **, *p* < 0.01; Bonferroni-corrected values).

**Figure 3.**
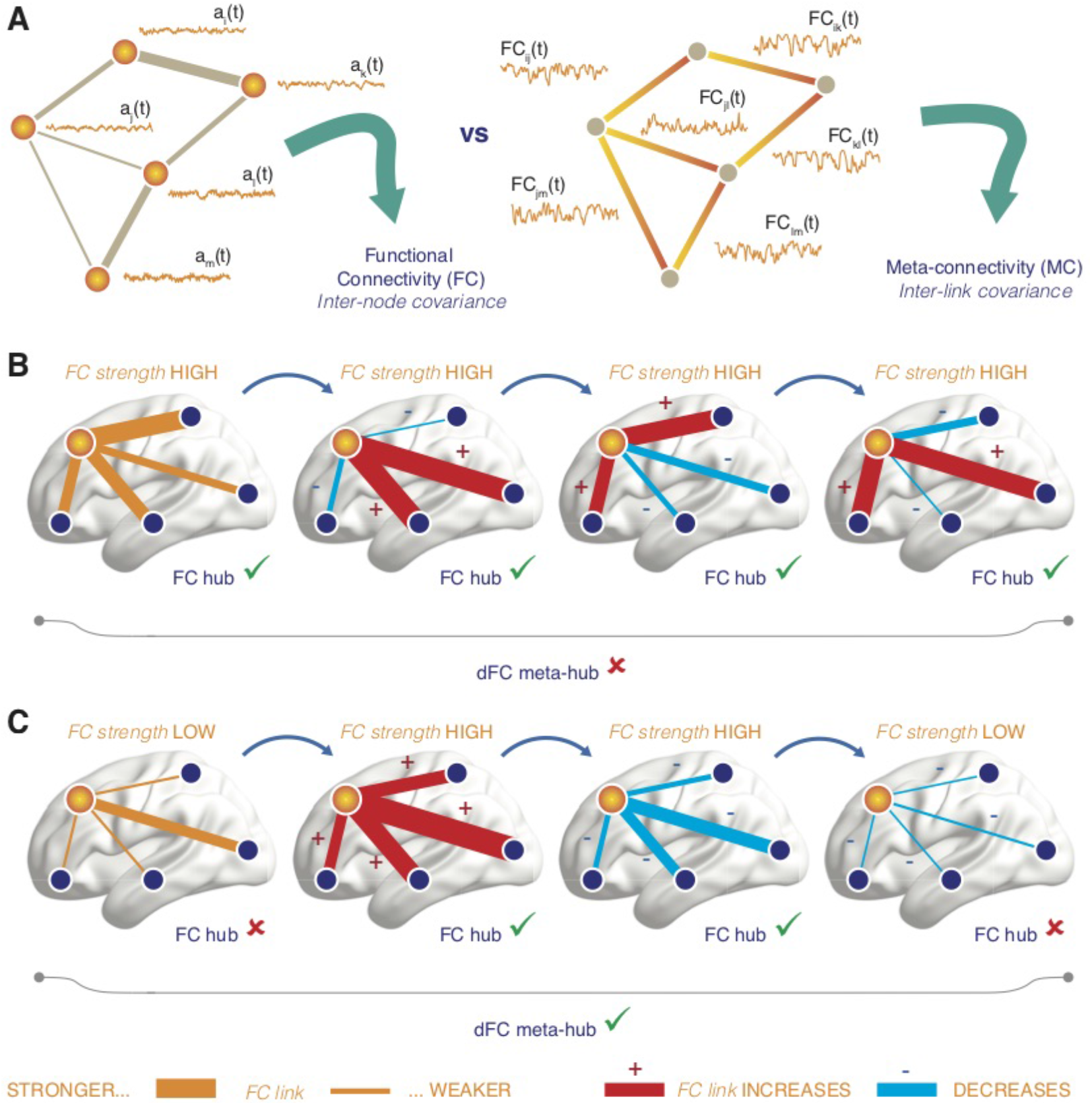
FC connectivity and hubs vs dFC meta-connectivity and meta-hubs. **A**. Functional connectivity describes correlations between fluctuations of network node activity, while *dFC metaconnectivity* (MC) describes correlations between fluctuations of network link time-resolved strengths. A node is said to be a *FC hub* if the fluctuations of its adjacent nodes are highly correlated. Analogously, a node is said to be a *dFC meta-hub* if the fluctuations of its incident links are highly correlated. The property of FC hubness can be assessed within every frame of a sequence of time-resolved FC(*t*) networks. Assessing the property of dFC meta-hubness requires instead to evaluate correlations between link fluctuations (compiled in the MC matrix based on an entire sequence of FC(*t*) frames. **B-C.** We show, for illustration, two short cartoon sequences of time-evolving FC subgraphs centered on a central node (in yellow). In the different network frames (time advancing from left to right) the thickness of a link is proportional to its instantaneous strength. FC links which increase their strength with respect to the previous frame are colored in red (and labeled by a small “+” sign), while FC links which decrease their strength are colored in blue (and labeled by a “-” sign). In panel **B** the sum of the weights of all links incident to the central node remain high at every time. Therefore the central node is a FC hub in all the shown time-frames, even if the fluctuations of its incident link have inconsistent directions. In panel **C**, in some of the frames the total strength of the central node is low, so that it is not anymore always a FC hub. However its incident links, independently on their instantaneous strength, change their strength in a consistent manner (they all increase, or decrease), allowing us to describe the central node as a dFC meta-hub (or coordination center of dFC).

**Figure 4.**
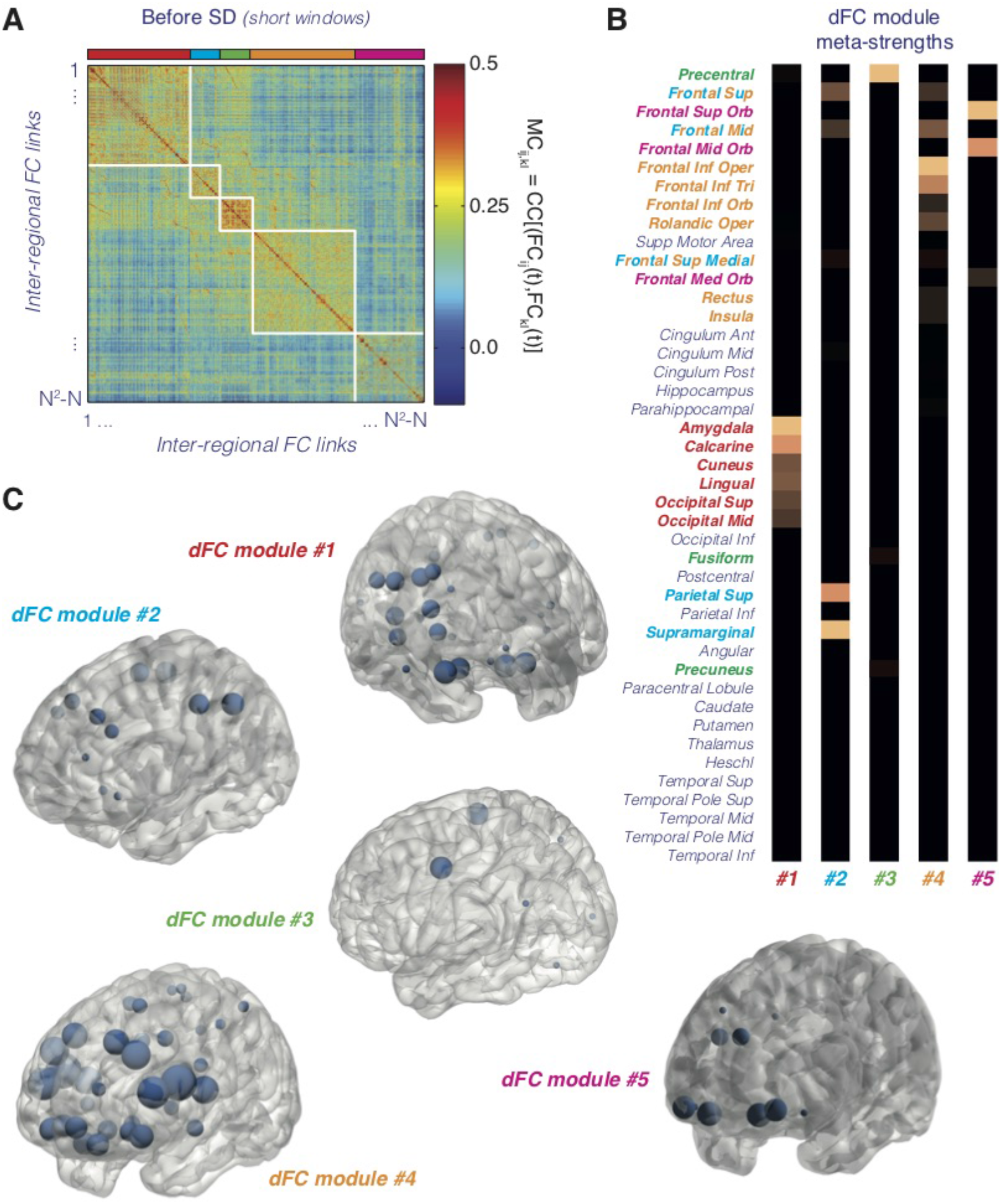
Meta-connectivity (MC) and dFC modules. **A.** Group-averaged MC matrix (see Figure 3A) for resting state fMRI sessions before SD (window-size of ~20 s; see Figure S4A for MC after SD). The order of the inter-regional FC links has been rearranged to emphasize the modular structure of the interlink correlation matrix. We have identified 5 different *dFC modules*, corresponding to independently fluctuating functional subnetworks. **B.** Summary plots of the meta-strengths of different brain regions restricted to the different dFC modules (symmetrized over the two hemispheres, given the lack of robust inter-hemispheric differences). The presence of a few large meta-strength entries for every dFC module indicates that different dFC modules are organized around distinct and specific sets of meta-hub regional nodes. **C.** Localization of the strongest meta-hubs within each of the 5 identified dFC modules. The color of the dFC module label is matched to the one of the associated strongest meta-strength regions in panel **B** (the region name is written with letters of multiple colors, when the region is a meta-hub for more than one dFC module simultaneously). Note that, although dFC modules have localized meta-hubs, they all influence nearly the whole brain (see Figure S4C).

#### Control resting state fMRI dataset

To verify that the key results of our analyses did not apply just to our specific dataset, we also repeated some of the analyses on an independent control dataset, selected from rs-fMRI data released as part of the Human Connectome Project (HCP), WU-Minn Consortium. We used the same selected subjects used for statistical benchmarking of functional network analyses in Termenon et al. (2016). This sample includes 100 young healthy adults aged 20 to 35 years (54 females). Each subject underwent two rs-fMRI acquisitions on different days and here, not being interested in test-retesting issue, we used only data from the first day sessions. For this sessions, TR = 720 ms and resting state scan duration was of 14 min and 24 s. The same parcellation as for the SD dataset was used. More details on data acquisition and pre-processing for this control dataset can be found on Termenon et al. (2016).

#### Extraction of time-dependent Functional Connectivity and dFC matrices

These methods have first been introduced by Battaglia et al. (2020) and we here provide a brief explanation. We estimated time-dependent Functional Connectivity matrices FC(*t*) by sliding a temporal window of fixed duration *W* (cf. Allen et al., 2012) and by evaluating zero-lag Pearson correlations between resting-state BOLD time series from different brain regions *i* and *j*:

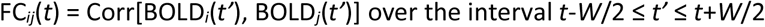

All entries were retained in the matrix, independently of whether the correlation values were significant or not or without fixing any threshold (i.e., we treated FC_ij_ entries as descriptive *features* operationally defined by the above formula).

To evaluate the dFC matrices of Figure 2 and S8 we introduced a notion of similarity between FC*(t)* matrices following (Hansen et al., 2015), based on the Pearson correlation between the entries of their upper-triangular parts:

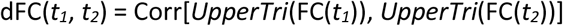

The dFC matrices thus depend on the window-size *W* adopted when evaluating the stream of time-dependent FC(*t*)’s. To perform the two-dimensional projections of the sequence of FC(*t*) matrices in Figure 2B we fed the vectors *UpperTri*(FC(*t*)) as input features into a t-Stochastic Neighborhood Embedding algorithm (exact method) as described by Hinton & Van der Maaten (2008), using default perplexity = 30 and exaggeration = 4 parameters in the used MATLAB^®^ (MathWorks R2017b) implementation.

#### Analysis of dFC speeds

Since we used correlation as a measure of similarity between matrices, we naturally chose to use the *correlation distance* between two observations of functional connectivity as a measure of the amount of change between two FC(*t*) observations. By measuring the distance between two FC observations separated by a fixed amount of time set to be equal to the window-size *W* we thus defined the instantaneous *global dFC speed* as:

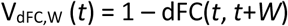

This dFC speed is termed “global” because evaluated in terms of comparisons of whole-brain FC matrices.

This definition makes dFC speed dependent on the chosen window size W. To improve the estimation of dFC speed histograms by increasing the number of sampled speed observations (not large for single window-size speed estimation, especially when *W* is large, given the need to consider not-overlapping windows), as well as for avoiding potential aliasing artefacts due to the use of a single window (Leonardi & Van de Ville, 2015), we decided to *pool* window sizes together, observing that speed distributions for close enough *W* were similar. More specifically, we realized that for a vast majority of subjects and for most dFC speed bins, binned counts in the histograms extracted at contiguous window sizes were statistically indistinguishable. Concretely, for 8 out of the 11 included subjects, the binned dFC speed counts in the histograms extracted at window sizes W_i_ and W_i+1_ were statistically indistinguishable (overlapping confidence intervals, Agresti-Coull estimation) for at least 14 out of 20 bins, where W_i_ are the window sizes included in our analyses (and the statement holds for any of the coinsidered W_i_’s, ranging from 3 to 30 TRs). Given this substantial degree of redundancy between speed distributions for contiguous window sizes, we chose to pool speed samples to form just two histograms, over two (arbitrary) window ranges: a “short” window-size range, 10 < *W* < 45 s; and a “long” window-size range, 45 < *W* < 80 s. Window pooling reduces the number of independent statistical comparisons to avoid the risk of false positive when comparing speeds before and after SD and leads to smooth dFC speed histograms (see also Battaglia et al. 2020). In the context of this study, however, we only focused on the median of pooled speed samples, evaluated separately for each of the two window-size ranges for each subject. For comparison, we nevertheless also show results also for single window analyses of global dFC speed prior to pooling in Figure S1.

As for cognitive performance variations, we evaluated the difference in global dFC speeds before and after SD: at the group level, using U-Mann-Whitney test; at the within-subject level, using paired Wilcoxon signed-rank test. Bonferroni correction was applied on the number of tested window-size ranges (ν = 2).

#### Meta-connectivity

Out of the stream of FC*(t)* matrices, we extracted *M* = *N*(*N*-1)/2 time series of pairwise FC couplings given by the entries FC_*ij*_*(t)* for all pairs of regions *i* and *j* with *i* < *j* ≤ *N*. We used a very short windowsize of *W* = ~19 s (7 TR), with a window step of 1 TR. The uncertainty in estimating FCs due to this unusually short *W* was compensated by the larger number of points in obtained FC coupling timeseries, which must be imperatively large in order to reliably compute inter-links correlations. The entries of the Meta-Connectivity matrix (MC) were then given by:

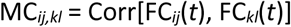

As for the FC analysis, we included all MC entries into the matrix, independently of whether the correlation values were significant or not or without fixing any threshold. We compiled all MC_*ij,kl*_ entries into a matrix format that allowed us to easily identify the pair of links involved into each meta-link and the participating brain regions. That is, we built MC matrices of (*N^2^-N*)-times-(*N^2^-N*) size, where different rows correspond to different directed pairs of regions – i.e. both the pair (*i,j*) and the pair (*j,i*) were included – and only links corresponding to self-loops – i.e. of the type (*i,i*) – were excluded. The MC representation described above had a substantial amount of redundance, since:

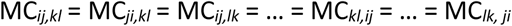

For the sake of computational efficiency we computed only *P* = *M*(*M*-1)/2 independent Pearson correlations between pairs of possible FC link time series FC_*ij*_(*t*) and FC_*kl*_(*t*), with *i* < *j, k* < *l, i* ≤ *k, j* < *l* and copied this inter-link correlation value into the eight degenerate MC matrix entries, in order to allow interpreting MC as an adjacency matrix between links.

We then defined the *meta-strength* of a node *i* as the sum of the meta-connectivity weights between all pairs of links incident on node *i*, i.e.:

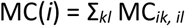

Besides the strengths MC_*ij,kl*_ of individual meta-links, we also computed integrated meta-connectivity strengths for individual links. A node with particularly large meta-strength is termed a *meta-hub*.

Note that MC matrices, evaluated for each single subject, can then be conveniently averaged over groups (analogously to FC matrices). The MC matrices shown in Figures 5A and S4A are thus averaged over the entire group of the 10 retained subjects.

**Figure 5.**
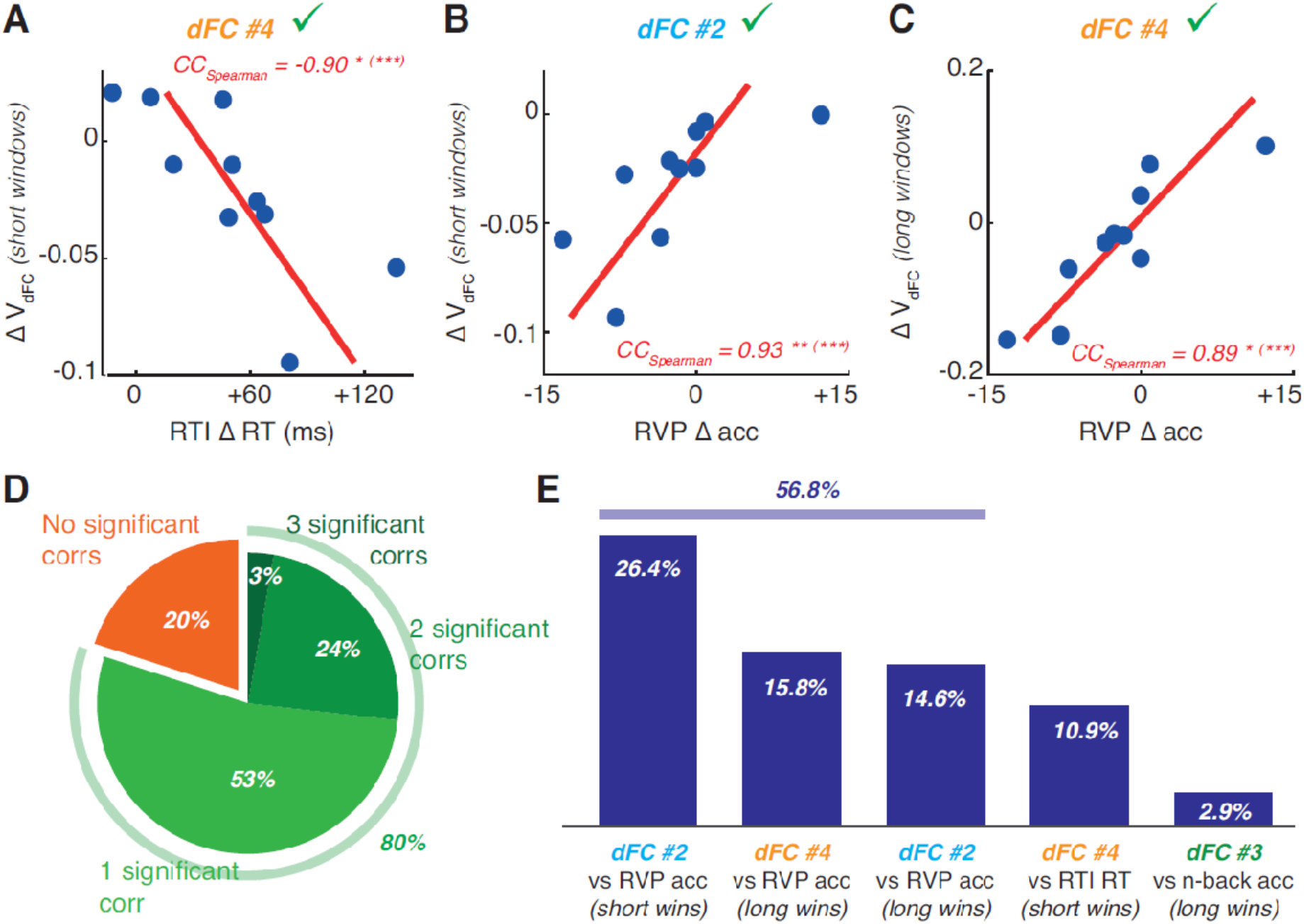
Modular dFC speed variations after SD correlate with variations of cognitive performance in specific tasks. Performing a separate dFC speed analysis for each of the identified dFC modules, we extracted *modular dFC speeds*. **A-C.** Selected scatter plots of subject-specific variations of cognitive performance scores and modular dFC speed. For the here represented decomposition into dFC modules, we highlight significant correlations: (**A**) between variations of RTI reaction time and modular speed of dFC module #4 on short window sizes; (**B**) between variations of RVP accuracy and modular speed of dFC module #2 on short window sizes; (**C**) between variations of RVP accuracy and modular speed of dFC module #4 on long window sizes (see Table 3 for other tendential correlations). D-E. The algorithm for extracting dFC modules from the MC matrix is stochastic and can yield different solutions, associated to potentially different patterns of correlation between modular dFC speed and cognition. Stars denote significance: *, *p* < 0.05; **, *p* < 0.01; ***, *p* < 0.001; Bonferroni-corrected values outside brackets (Bonferroni-uncorrected within brackets). D. Among 2000 different instances of modular decomposition, 80% displayed at least one robust correlation between variations of modular dFC speed and cognitive scores. E. The most robust correlations were the ones between fronto-parietal and parietal modules dFC #2 and dFC #4, arising in 60% of cases, in addition to other eventual secondary correlations. See S9 for additional analyses on dFC modular speeds.

#### Edge Functional Connectivity

For comparison with MC analyses, we also computed edge-centric Functional Connectivity (eFC) as introduced by Faskowitz et al. (2019). Similarly to MC, eFC equally provides a description of correlations between link fluctuations, however eFC differs from MC in the way in which the temporal stream of dFC is estimated. While in MC we rely on a sliding window approach to extract FC(*t*), in the case of eFC windowing is not used but signal correlation fluctuations are studied at the instantaneous level. The time-series BOLD_*i*_(*t*) are first z-scored, then time-series of pairwise products are constructed:

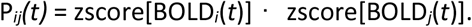

Finally, eFC is constructed as the correlation matrix of these instantaneous pairwise product time-series:

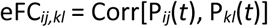

#### Extraction of dFC modules

Packaged in the redundant format described, the MC matrix can be considered as the adjacency matrix of a graph whose nodes are FC links. Thus, communities of temporally co-varying FC links can simply be evaluated by applying any algorithm for node-community detection to the MC matrix. We thus extracted modules by applying a standard Louvain algorithm on the group-averaged MC matrix (before SD), adapted to weighed undirected graphs with mixed positive and negative entries (“symmetric treatment” in the Louvain method’s implementation in the Brain Connectivity Toolbox by Rubinov & Sporns (2010)). The Louvain method has one adjustable parameter whose default value is Γ = 1. By taking larger (or smaller) values, the algorithm will tend to return modular decompositions with a larger (smaller) number of modules. We chose to use Γ = 1.045 at the center of a range reliably and robustly returning 5 modules. Larger values of Γ were on the contrary leading to modular decompositions with fluctuating numbers of modules.

Even when Γ is fixed the algorithm still returns different modular decompositions every time it is run, because it involves a stochastic optimization procedure. We thus generated 2000 alternative modular decompositions to study how variable or robust they were. To compare two modular decompositions M_1_ and M_2_ we first identified homologous modules, by relabeling the modules in order to have minimum Hamming distance – i.e. number of different symbols in an ordered sequence – between the module label sequences for M_1_ and M_2_. Then we computed the meta-strengths of every region module by module – i.e. restricting the MC matrix to include only rows and columns corresponding to links belonging to the considered module – for both modular decompositions M_1_ and M_2_. Finally, we correlated the sequence of meta-strengths for equally-labelled modules in M_1_ and M_2_, to assess how similar modules were in terms of their meta-hubs and node ranking in terms of meta-strengths. The large inter-decomposition meta-strength correlations reported in S9A for all modules confirm that our choice of decomposing MC into 5 modules leads to robustly preserved modules (and their associated meta-hubs).

To construct Figure S5E, we extracted 500 alternative modular decompositions of the MC matrix mediated from the control HCP dataset and maximally aligned each of them to the reference modular decomposition of Figure 4A. For each of these realigned modular partitions a confusion matrix with the reference modular decomposition was computed and then averaged this confusion matrix over the 500 alternative modular structures.

#### Modular and RSN-restricted dFC speed analyses

A dFC module is a collection of functional links and is, therefore, describing a sub-graph of the temporal network (Holme & Saramäki, 2012) formed by the stream of whole-brain FC(*t*). We denote as FC^α^(*t*) the stream of sub-graphs obtained keeping in the stream FC(*t*) only the links included in the considered dFC module #α. It becomes thus possible to compute *modular* dFC matrices or dFC speeds by adopting precisely the same formulas as for global dFC matrices and speeds, but replacing FC(*t*) with FC^α^(*t*).

For comparison, we also restricted dFC analyses to sub-graphs made by all the links between regional nodes assigned to a reference Resting State Network (RSN). For mapping the 86 regions of the AAL atlas adopted here to the Yeo7 atlas, including 8 RSNs –visual (V), somato-motor (SM), Dorsal and Ventral Attention (A), Fronto-temporal (FT), Fronto-parietal (FP), Default mode (DM) and Subcortical (SC)–, we used the correspondence introduced by Amico et al. (2017). Given that the Dorsal Attentional Network would include just one link under this mapping, preventing evaluation of correlation distance between frames in the dFC network, we merged it with the Ventral Attentional Network into a common Attentional RSN.

As for global dFC speeds and cognition, we evaluated difference in modular or RSN-restricted dFC speeds before and after SD at the within-subject level (S8B-C), using paired Wilcoxon signed-rank test. Bonferroni correction was applied on the number of tested window-size ranges and dFC modules (ν = 2 x 5 = 10) or RSN networks (ν = 2 x 7 = 14).

#### Correlations between variations of dFC speed and of cognitive performance

Given the large inter-subject variability of both dFC speed and cognitive performance, we chose to correlate the *changes* of these variables, i.e. the difference or dFC speeds (or cognitive scores) measured after and before SD. We then computed Spearman correlation between changes in dFC speed and changes in cognitive scores. To assess significance of these correlations we applied a conservative Bonferroni correction to the *p*-values of these correlations. For correlations with global dFC speed, we corrected on the number of window ranges and the number of cognitive scores (ν = 2 x 5 = 10). For correlation with modular dFC speeds, we further corrected for the number of modules (ν = 2 x 5 x 5 = 50).

#### Correlation between variations of dFC speed and EEG

Only in 4 subjects out of 11, sleep stage N3 was reached (and only in fMRI sessions after sleep deprivation). All epochs in sleep stage N3 were rejected from fMRI analyses. For 3 of the 4 subjects reaching stage N3 these epochs represented less than 20% of the total scan duration. For the 4^th^ subject, the duration of sleep instage N3 reached 80% of scan duration, therefore we had to reject this subject for all within-subject analyses of dFC variations between before and after sleep deprivation. We checked that dFC fluctuations were not trivially due to switching between awake, N1 and N2 states. As shown by Figure S2, genuine BOLD dFC fluctuations were observed in all three brain states (not surprisingly, since intrinsic cognitive activities are not suppressed during sleep), with overall mild differences of speed. Most likely, the lack of significance of dFC speed comparison between sleep stages over most window sizes was due to the small amount of observations of speed available when considering sub-segments of already relatively short fMRI scans. Therefore for the other analyses of the study and to achieve superior statistical power, we pooled together speed observations in awake (overall 69% of analyzed fMRI time points), N1 (overall 13% of analyzed fMRI time points) and N2 (18%) epochs. This prevented studying differences of dFC speed within scan epochs but allowed a better inter-scan comparison.

#### dFCwalk toolbox

We have compiled a toolbox of MATLAB^®^ functions to perform most dFC and MC analysis operations described in the present study. It can be downloaded as Support File Sx and will be described in a *MethodsX* paper in preparation.

## Results

### Sleep deprivation heterogeneously affects cognitive performance

We first considered how performance in different cognitive tasks is affected by 24 h of SD. We studied performance on three different tests: a Reaction Time test (RTI), providing a basic assessment of the speed of motor response; an n-back task, assessing working memory using sequences of letters (n-back); and a Rapid Visual Processing test (RVP), providing a measure of sustained visual attention (see *Materials and Methods* for details).

The variations in cognitive performance induced by SD which we observed at both the group and the single-subject levels are summarized in Table 1. Relative variations observed at the single-subject level are also graphically shown in Figure 1. We found that cognitive change induced by SD was highly heterogeneous. While for all three tests, reaction time tended to increase and accuracy to decrease, the actual variations could differ from up to two orders of magnitude across the different subjects. As a result, none of the comparisons at the group level between performance indicators before and after SD remained significant after applying multiple comparison correction (Table 1, left columns).

**Table 1.**
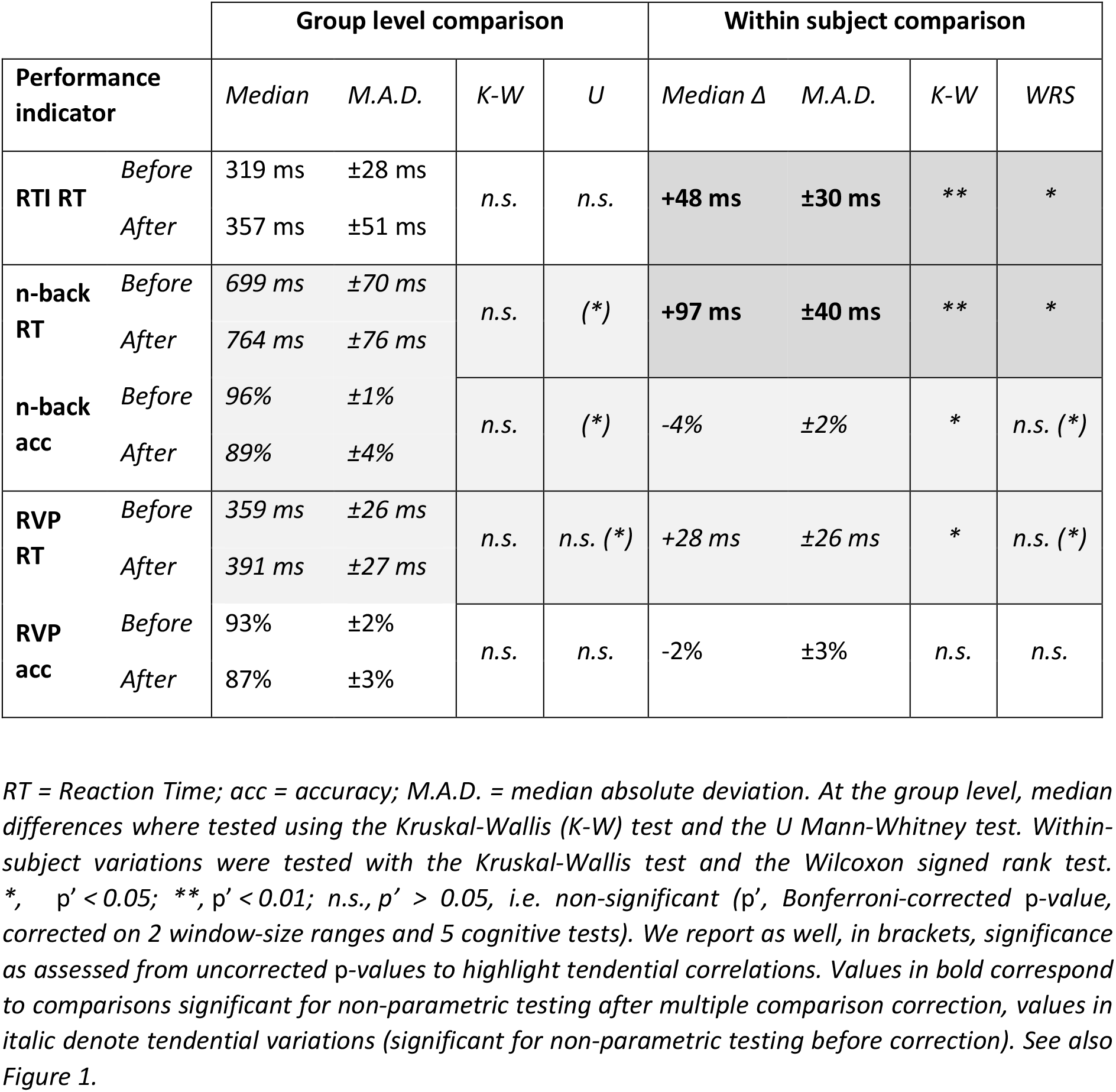
Variations of cognitive performance induced by sleep deprivation.

|  |  | Group level comparison |  |  |  | Within subject comparison |  |  |  |
| --- | --- | --- | --- | --- | --- | --- | --- | --- | --- |
| Performance indicator | | Median | M.A.D. | K-W | U | Median $\Delta$ | M.A.D. | K-W | WRS |
| **RTI RT** | *Before* | 319 ms | $\pm 28$ ms | *n.s.* | *n.s.* | **+48 ms** | **$\pm 30$ ms** | **\*\*** | **\*** |
| | *After* | 357 ms | $\pm 51$ ms | | | | | | |
| **n-back RT** | *Before* | 699 ms | $\pm 70$ ms | *n.s.* | **(\*)** | **+97 ms** | **$\pm 40$ ms** | **\*\*** | **\*** |
| | *After* | 764 ms | $\pm 76$ ms | | | | | | |
| **n-back acc** | *Before* | 96% | $\pm 1\%$ | *n.s.* | **(\*)** | -4% | $\pm 2\%$ | **\*** | *n.s. (\*)* |
| | *After* | 89% | $\pm 4\%$ | | | | | | |
| **RVP RT** | *Before* | 359 ms | $\pm 26$ ms | *n.s.* | *n.s. (\*)* | +28 ms | $\pm 26$ ms | **\*** | *n.s. (\*)* |
| | *After* | 391 ms | $\pm 27$ ms | | | | | | |
| **RVP acc** | *Before* | 93% | $\pm 2\%$ | *n.s.* | *n.s.* | -2% | $\pm 3\%$ | *n.s.* | *n.s.* |
| | *After* | 87% | $\pm 3\%$ | | | | | | |

RT = Reaction Time; acc = accuracy; M.A.D. = median absolute deviation. At the group level, median differences were tested using the Kruskal-Wallis (K-W) test and the U Mann-Whitney test. Within-subject variations were tested with the Kruskal-Wallis test and the Wilcoxon signed rank test. \*, $p' < 0.05$ ; \*\*, $p' < 0.01$ ; *n.s.*, $p' > 0.05$ , i.e. non-significant ( $p'$ , Bonferroni-corrected p-value, corrected on 2 window-size ranges and 5 cognitive tests). We report as well, in brackets, significance as assessed from uncorrected p-values to highlight tendential correlations. Values in bold correspond to comparisons significant for non-parametric testing after multiple comparison correction, values in *italic* denote tendential variations (significant for non-parametric testing before correction). See also Figure 1.

However, when considering the variation of performance within every single subject (Table 1, right columns), we found that its direction was more consistent across subjects. As a result, all median within-subject variations of reaction time were significantly positive (Kruskal-Wallis testing of medians; however, only RTI and n-back RT variations were significant under the more conservative Wilcoxon Signed Rank test). Furthermore, the median within-subject variation of n-back accuracy was also significantly negative. On the contrary, for RVP accuracy the results were more variable and some subjects even increased their RVP accuracy after SD.

We can thus confirm that sleep deprivation affects cognitive performance in all the considered tests. However, the effects of SD were highly heterogeneous and often small, thus providing an interesting benchmark for our attempt of predicting variations of cognitive performance with dFC speed metrics.

### Sleep deprivation slows down the speed of global dFC

We then studied how dFC is affected by SD at the whole brain level. We focused in particular on quantifying the rate of reconfiguration of brain-wide FC networks during resting state and its variations after 24 h of sleep deprivation.

To estimate time-resolved FC networks, we relied on a simple sliding window approach (Allen et al., 2012). We considered the whole time series of the BOLD signal for each of the regions of the parcellation, then divided the time series in non-overlapping windows of size *W*, computing a different FC network FC(*t*) in each of these windows (Figure 2A). To quantify the amount of FC network variation from one time window to the next, we calculated the pairwise correlation between subsequent functional connectivity matrices and evaluated instantaneous *global dFC speed* at time *t* as *V_dFC, w_*(*t*) = 1 – Corr[FC(*t*), FC(*t+W*)], where Corr denotes the Pearson Correlation Coefficient between the weighed FC connectivity matrix entries (see *Materials and Methods*). In other words, the more (the less) the FC network at a given time is correlated with FC after a waiting time equal to the window size *W*, the smaller (the larger) the estimated global dFC speed will be at this time.

The obtained values of global dFC speed thus necessarily depend on the chosen window size *W*. However, the obtained counts of dFC speed in binned histograms can hardly be statistically distinguished (see *Materials and Methods*) when neighbouring window sizes are considered. We therefore chose to refine the estimation of the median speed by pooling dFC speed estimates computed for multiple contiguous window sizes together (see Battaglia et al., *companion manuscript* and *Materials and Methods*). Pooling different windows (with incommensurate sizes) furthermore protects against potential aliasing artifacts (Leonardi & Van de Ville, 2015) deriving from the adoption of unique fixed window size (see *Discussion*). In this study, we chose to pool single-window dFC estimates into two different ranges, separately estimating dFC speeds for *“short”* (10 to 40 s) and *long* (40 to 80 s) window lengths. The chosen window lengths included very short windows that are not usually adopted when estimating FC. See *Discussion* and, especially, the companion study by Battaglia et al. (2020) for a thorough discussion on the rationale behind this choice.

As previously mentioned, in contrast with other approaches like clustering (see e.g. Allen et al., 2012) or Hidden Markov Modeling (Vidaurre et al., 2016), we do not extract “states” of functional connectivity in our dFC speed approach. Instead, we quantify the speed of transition between sequential ‘snapshots’ of functional connectivity along a random walk in the high-dimensional space of FC. The speed of reconfiguration is not constant along this random walk, but there is a stochastic alternation between epochs of time in which dFC speed slows down – dFC “knots” –, and other epochs in which dFC speeds-up – or “leaps” –, suggesting that the sampling of functional connectivity space is not a Brownian-like motion – like a particle of dust floating in the air – but rather a realization of an anomalous stochastic process – e.g. like the fast-spreading of epidemics in the modern world or animals optimally foraging for food – (see Battaglia et al., 2020, for a thorough discussion).

In Figure 2B, we provide two visualizations of these knots and leaps along resting-state dFC for a representative subject. In the top row, we show the so-called *dFC matrices* (Hansen et al., 2015) associated to the two resting sessions, performed first before (left) and then after SD (right). In these matrices, the entry dFC(*t_1_, t_2_*) provides the correlation between the network FC(*t_1_*) observed at a time *t_1_* and a second network FC(*t_2_*) observed at a second time *t_2_*. In such dFC matrix representations, dFC knots appear as red-hued (high correlation, hence low speed) squares along the diagonal and dFC leaps as crossing blue stripes (low correlation, hence high speed). Even at visual inspection, dFC knots are markedly evident in the dFC matrix extracted after SD.

In the bottom row of Figure 2B, we show dimensionally-reduced representations of the random walk in FC network space associated with the same sessions imaged in the dFC matrices plotted above. We obtained such two-dimensional projections by adopting a powerful non-linear dimensional reduction technique, t-Stochastic Neighborhood Embedding (t-SNE, Hinton & van der Maaten, 2008). In these plots, each dot corresponds to the projection of a different time-resolved FC network and temporally consecutive dots are linked by a line (see *Materials and Methods*). The dFC random walk before SD appears relatively smooth and continuous, interrupted only by a few cusp points, both before and after SD.

Beyond the visualizations of Figure 2B, we then quantify the variations of global dFC speed before and after SD (Figures 2C and 2D). Figure 2C represents a boxplot of the distributions of global dFC speed evaluated for pooled short (right) and long (left) time windows. As discussed extensively in Battaglia et al. (2020), these distributions are generally skewed and non-Gaussian (i.e. associated to non Brownian dFC random walks). For both short and long window sizes, we found, on the group-level, that global dFC speed slowed down after 24 hs of sleep deprivation (U Mann-Whitney, Bonferroni-corrected *p*-values: *p* = 0.025 for short windows; *p* = 0.014 for long windows). Note that we would have obtained the same result if global dFC speed for individual window sizes had been computed without pooling (Figure S1). Analogously, Figure 2D shows a boxplot of within-subject variations, which were also significantly negative (Wilcoxon Signed Rank, Bonferroni-corrected *p*-values: *p* = 0.004 for short windows; *p* = 0.02 for long windows). The effect of sleep deprivation on global dFC speed is thus very robust, slowing occurring also on the subject by subject level – despite the heterogeneity of baseline speeds before SD – and not just at the group median level.

We finally checked whether the fluctuations of dFC speed along the resting state sessions were related to the occurrence of short sleep episodes (detected via simultaneous EEG while in the MRI scanner). EEG data was manually sleep-staged for consecutive 30 s blocks by an expert (Awake, N1, N2, N3, according to the AASM sleep scoring scheme, http://www.aasmnet.org/). Although some of the subjects displayed transient sleep episodes, especially after the stress of SD (see *Materials and Methods* for subject inclusion and rejection criteria), we observed that dFC network reconfigurations did not stop during sleep but were continually ongoing at a non-vanishing speed. In particular – most likely because of the limited amount of data – we could not find a systematic impact of transiently entering into sleep stages and the concurrently measured dFC speeds. In Figure S2A, we show that median dFC speeds tended to slow down at the single-subject level during sleep stage N1 with respect to awake state, however, this effect was only tendentially significant for a few window-sizes belonging to the long window sizes range. When considering on the contrary the deeper sleep stage N2, significant slowing of median dFC with respect to wake could not anymore be detected (Figure S2B). Therefore, the variations of global dFC speed, especially for short window sizes, cannot be explained merely in terms of an increased rate of occurrence of transient sleep episodes inside the scanner.

### Global dFC speed variation does not correlate with cognitive performance

Both cognitive performance and global dFC speed were affected by SD. We therefore assessed whether the cognitive change induced by SD correlated with the monitored dFC speed reductions within each subject. However, as compiled in Table 2 (left columns), for short window sizes, global dFC speed reductions did not significantly correlate with variations in reaction time or accuracy on RTI, n-back or RVP. When considering the long window sizes, none of the correlations between the slowing of global dFC speed and cognitive performance survived multiple comparison corrections (Table 2, right columns). Figure S3 shows scatter plots revealing lack of correlation (for short window sizes, Figure S3A-C) and (for long window sizes, Figure S3D-F) between global dFC speed and cognitive performance variation in the three considered tasks.

**Table 2.** Spearman correlations between within-subject variations of global dFC speed and cognitive change after SD.

| | Short window sizes $\Delta V_{dFC}$ | | Long window sizes $\Delta V_{dFC}$ | |
| --- | --- | --- | --- | --- |
|  | *Correlation* | *p* | *Correlation* | *p* |
| **RTI $\Delta$ RT** | -0.58 | n.s. | *-0.71* | *0.028 (\*)* |
| **n-back $\Delta$ RT** | -0.29 | n.s. | *-0.76* | *0.037 (\*)* |
| **n-back $\Delta$ acc** | 0.63 | n.s. | *0.75* | *0.025 (\*)* |
| **RVP $\Delta$ RT** | 0.03 | n.s. | -0.43 | n.s. |
| **RVP $\Delta$ acc** | 0.44 | n.s. | *0.79* | *0.006 (\*\*)* |

*RT = Reaction Time; acc = accuracy. Spearman correlation values between within-subject variations of global dFC speeds and within-subject variations of cognitive performance scores. We report the associated uncorrected p-values, as well as significance. \*, $p' < 0.05$ ; \*\*, $p' < 0.01$ ; n.s., $p > 0.05$ , i.e. non-significant ( $p'$ , Bonferroni-corrected p-value, corrected on 2 window size ranges and 5 cognitive tests). We also report (in brackets), significance as assessed from uncorrected p-values to highlight tendential correlations (correlation values in *italic* and light shaded table entry are significant before correction, i.e. (\*), $p < 0.05$ ; (\*\*), $p < 0.01$ ). See also Figure S3.*

Overall, we thus found weak evidence for consistent correlations between global dFC slowing and cognitive performance and reaction times. This result is in apparent discrepancy with the positive correlation found between global dFC study and a general cognitive performance score (MOCA; Nasreddine et al., 2005) in the companion study by Battaglia et al. (2020). While it is possible that this result is due to the small number of subjects in our study (see *Discussion*), we also hypothesized that it may reflect the fact that changes in dFC relevant to specific cognitive processes occur uniquely within restricted sub-networks which are relevant for the considered tasks. Therefore, these task-related dFC alterations could be buried under the fluctuations of many other irrelevant networks, when computing global speed by averaging dFC over the whole brain. In order to confirm our hypothesis, we introduced a novel approach, which allows focusing on dFC fluctuations of specific brain sub-networks and their correlations with cognition.

### Beyond global dFC: meta-connectivity, dFC modules and meta-hubs

The fluctuations of FC networks giving rise to dFC can be decomposed into the fluctuations of individual pairwise inter-regional correlations. A fluctuating network can thus be seen as a collection of fluctuating links (Figure 3A, to be compared with Figure 2A). It thus becomes logical to look for groups of links (i.e. sub-graphs of the whole brain FC graph) whose fluctuations in time are more correlated between each other than they are with the fluctuations of other groups of links. In order to search for such coherently fluctuating sub-networks, we computed a correlation matrix between link-fluctuations (Davison et al., 2015; Brovelli et al., 2017; Faskowitz et al., 2019) – which we here call *meta-connectivity (MC) matrix* – and applied to this matrix a standard community detection algorithm in order to extract modules of covarying links (*dFC modules*).

As a matter of fact –and as discussed as well in Faskowitz et al. (2019), where it is referred to as “edge functional connectivity” (eFC)– MC provides a static representation of the spatial organization of dynamic fluctuations of FC links overtime. In this sense, it constitutes a generalization of the classical notion of Functional Connectivity from nodes to links. The FC matrix provides indeed a static representation of the dynamic fluctuations of regional nodes along time. Strong FC between two nodes indicates that their fluctuations in time are strongly correlated. Analogously, strong MC between two links indicates that their fluctuations in time are strongly correlated. MC analysis thus captures the fact that the fluctuations of different links are not independent but are, on the contrary, functionally coupled. The static MC matrix epitomizes the underlying statistical coordination structure that, any moment in time, constrains the generation of interdependent link fluctuations.

In conventional FC analyses, one often defines regional *hubs*, as regional nodes such as the fluctuations of their *adjacent* nodes are strongly correlated between them. Note that two nodes in a graph are commonly defined as “adjacent” if connected by a link (Bollobas, 1998). The definition of hubs allows identifying coordination centers in the FC network, which are clearly anatomically localized and may correspond to specific functional roles in cognition (Bullmore & Sporns, 2009; Sporns & Betzel, 2016). When considering, however, dFC and MC, it is more intricate to generalize this notion of hub, because MC is a link-based covariance and a functional link is more widely distributed and less “localized” in the cortex, since it spans over distances. We here nevertheless introduce the notion of *meta-hub*, as regional nodes such as the fluctuations of their *incident* links are strongly correlated between them. Note that a link in a graph is commonly defined as “incident” to a node if attached to this node (Bollobas, 1998). Meta-hubs can thus be seen as control centers coordinating the fluctuations of the links originating from them. Adopting a graph-theoretical jargon, we can equivalently define meta-hubs as the centers of *star FC sub-graphs* (Bollobas, 1998) whose links – the rays originating from the center of the star – are strongly meta-connected. As we will see in the next section, each of the retrieved dFC modules is dominated by a small number of strongly meta-connected stars centered on a few module-specific meta-hubs (Figure 4).

In Figures 4B and 4C, we further elaborate on this notion of meta-hub and its distinction from the conventional notion of hub by providing a few cartoon examples. Both these figure panels display sequences of snapshots of a fluctuating star sub-network centered on a node of interest. In the sequence of Figure 3B, the *strength* of the central node – i.e. the overall sum of the weights of all it’s incident links – is steadily large. Correspondingly, this center node is a FC hub in all the considered snapshots. However, the different links incident to it fluctuate in an incoherent manner, some are strong when the others are weak, etc. Therefore, in this example, the central node is a hub with a large FC strength but *not* a meta-hub. Conversely, in Figure 3C, the strength of the central node fluctuates in time, since sometimes strong and sometimes weak. This central node is thus a hub only intermittently (and may not appear as such when considering connectivity averaged along time, as in static FC analyses). However, when an incident link is strong (or weak), then all the other incident links are strong (or weak) as well. In other words, link fluctuations are correlated – i.e. the plotted links are meta-connected – and the center node serves as a meta-hub, with a large *dFC meta-strength*, defined as the sum of the meta-connectivity between all its incident link pairs (see *Materials and Methods*).

### Resting-state dFC modules

In order to infer dFC modules, we evaluated the MC matrix before and after SD for each subject and for different window sizes. For MC estimation, we considered only short window sizes to avoid generating exceedingly smoothed time-series of dynamic FC weights with too few independent observations (see *Discussion*). We then averaged MC matrices obtained for different subjects and window sizes to obtain a common reference MC and finally ran a community detection algorithm in order to identify dFC modules (see *Materials and Methods* for details of the procedure).

Figure 4A shows this reference MC matrix for all sessions before SD. The MC matrix averaged over sessions after SD is shown in Figure S4A. As evident from the visual comparison of the two MC matrices before and after SD and also from the high linear correlation between MC entries (Pearson correlation 0.97, p < 0.001; see linear fit in Figure S4B), the modular structure is well preserved across the two conditions. For further comparison, we also computed the edge Functional Connectivity (eFC) as defined by Faskowitz et al. (2019) (see Materials and Methods). Once again, the ensemble averaged eFC matrix shown in Figure S5A displays a very similar modular structure as the MC matrix of Figure 4A. There is furthermore a very strong correlation between MC and eFC, down to the level of individual matrix entries (Figure S5B). Given this large degree of similarity, in this study, we, therefore, ignore these small differences and always use the modules extracted from the averaged MC matrix before SD of Figure 4A as reference modules. dFC modules can be clearly identified in the reference modular decomposition here considered (see below for analyses of the robustness of these modules).

All the modules include links that are distributed widely through the entire brain. Figure S4C shows which regions are incident to at least one link included in each of the five modules. Such representations reveal that dFC modules are highly overlapping in their regional reach and are essentially not-localized networks. As a matter of fact, Figure S4C reveals that all the modules have a “reach”, i.e. a set of regions on which they are exerting an influence, which is very close to… essentially the whole brain! In this sense, therefore, they are fundamentally different from conventional FC modules, which are internally integrated functional networks, but segregated between them. Here, every region is on the contrary “touched”, influenced by most dFC modules (see *Discussion*).

Nevertheless, every dFC module is organized around a specific set of localized regional controller nodes. When considering links which are strongly meta-connected (i.e. the more correlated), we found that each of the dFC modules includes prominent star sub-graphs which are centered on specific meta-hubs (cf. Figure 3C), different for each of the dFC modules. Figure 4B shows indeed the total meta-strengths of different brain regions within each of the five dFC meta-modules. It is clear that meta-strengths are not homogeneous but that each dFC module contains only a few meta-hubs with higher meta-strengths. In Figures 5C, we visualize the localization of these meta-hubs for each of the five modules.

A first dFC module (dFC #1) was organized around meta-hubs located bilaterally (i.e. always both in left and right hemisphere) in the amygdala and several regions involved in visual sensory processing, such as occipital or posterior regions, as calcarine sulcus, cuneus and superior and middle occipital gyrus.

A second dFC module (dFC #2) included a combination of parietal meta-hubs, such as supramarginal gyrus and left and right superior parietal lobule, and frontal meta-hubs such as superior, medial and middle frontal gyri.

A third dFC module (dFC #3) was organized around highly dominant meta-hubs in precentral gyrus, fusiform gyrus and precuneus.

A fourth module (dFC #4) was mainly dominated by meta-hubs situated in inferior frontal gyri (pars opercularis, pars triangularis and pars orbitalis), rolandic operculum, rectus gyrus and insulae as well as, shared with the second dFC module, superior, medial and middle frontal gyrus.

Finally, a fifth module (dFC #5) was centered on meta-hubs, again, within superior frontal gyrus, superior, middle and inferior frontal gyri (orbital parts) as well as medial orbitofrontal cortices.

Importantly, we could confirm the existence of an analogous modular structure in the resting state MC matrix extracted from a completely independent control dataset (see *Materials and Methods*). In Figure S5C, we show a resting MC averaged over 100 cognitively normal subjects mediated from the Human Connectome project (see Termenon et al., 2016), displaying a block community structure very similar to the one of Figures 4A and S4A). As shown by Figure S5D, the meta-strengths of this HCP-derived resting state MC of Figure S5C had a correlation of 0.99 (bootstrap 95% c.i., 0.993 < CC < 0.996) with the meta-strengths of the reference MC of Figure 4A. Furthermore, the confusion matrix of Figure S5E, confirms quantitatively that the modules extracted from the HCP-mediated MC had a substantial degree of overlap with the ones displayed in Figure 4B.

We note that these dFC modules only have a loose correspondence with modules that could be defined at the level of static FC. We show, in Figure S6A, static FC before sleep deprivation, averaged over the same group of subjects considered to extract the meta-connectivity of Figure 4A. We then extracted five modules even from this matrix, using the same unsupervised community detection algorithm (see *Materials and Methods*) and correlated the strengths of regional nodes within the resulting FC modules with the regional meta-strengths within the five dFC modules. The aim of this unsupervised analysis was not to extract static FC subnetworks –for this endeavor rather see e.g. Yeo et al. (2011)– but rather to check whether how much of the modular structure found at the level of MC was trivially redundant with information that could have been extracted with simpler analyses at the level of FC. As shown in Figure S6B, the dFC modules #1 and #5 could be put in correspondence with associated FC modules, with meta-hubs of dFC overlapping at a large extent with hubs of FC (meta-strength vs strength correlations for matched dFC and FC modules were *Corr* = 0.58 for dFC #1 and *Corr* = 0.59 for dFC #5, *p* < 0.001 after Bonferroni correction). However, for other modules, the correspondence was much less clear. Indeed the dFC #2 and #dFC #4 module both had a partial overlap with a same FC module (i.e. they could not be separated at the level of static FC analysis) and the dFC module #1 did not significantly overlap with any of the FC modules.

We also compared the localization of meta-hubs for our 5 dFC modules with a partition in standard Resting State Networks (RSNs), according to the Yeo7 atlas (Yeo et al., 2011; Amico et al., 2017; see *Materials and Methods* for details of the mapping). As revealed by Figure S7A, meta-hubs for the different dFC modules do not perfectly align with classic RSNs. The meta-hubs for the dFC module #1 has a good overlap with the visual RSN and meta-hubs for dFC module #2 have an imperfect overlap with a combination of the Dorsal and ventral attention RSNs, but some other RSNs (or dFC modules) do not have a clear counterpart.

Therefore, our unsupervised modular MC analyses bear some genuinely new information with respect to static FC analyses or partition in classic RSNs.

### Modular dFC speed variations are heterogeneous across modules

Each of the dFC modules can be considered, by definition, as a collection of functional links whose fluctuations are relatively independent from the fluctuations of links within other dFC modules. As a consequence, it is logical to perform independent dFC speed analyses restrained to links belonging to each of the dFC modules separately. For each dFC module, we thus extracted different *modular dFC matrices* and distributions of *modular dFC speed*. S8A shows examples of modular dFC matrices for a representative subject. Even at visual inspection, it appears that the red squares associated to dFC knots are generally not overlapping in time between the dFC modules. This result was expected since these modules have been separated precisely because of the relative independence of their temporal fluctuations. S8B-C shows boxplots of the within-subject variations of modular dFC speeds after sleep deprivation, for long (S8B) and short (S8C) time windows. We observed for all five dFC modules and for both ranges of window sizes a general tendency to a slowing of modular dFC speed, paralleling the result already found at the level of global dFC speed (cf. Figure 2D). However, this within-subject slowing of modular dFC speed could be proven to be significant only for the dFC #5 module, for both short and long time-windows and for the dFC #1 module, only for short time windows (see S8B-C). The non-uniformity across the dFC modules of speed variations confirms once again their relative degree of dynamic independence.

We also computed dFC speed restricted to subnetworks composed of all links between regional nodes assigned to each of the different RSNs, under the mapping to Yeo7 atlas (see *Materials and Methods*). As visible in Figures S7B and S7C, respectively for long and short window sizes ranges, also these RSN-restricted estimations of dFC speed show a heterogeneous tendency to slow down after SD. However, none of these RSN-restricted dFC speed reductions remained significant after corrected for multiple comparisons.

### Modular dFC speed variations correlate with cognitive performance variations

We then finally assessed if the variations of modular dFC speed after SD correlated with cognitive performance. Modular dFC speed variations displayed overall richer and more variegated patterns of correlations with cognitive performance than global dFC speed variations (see *Discussion*). Significant correlations were found with RVP accuracy, in a particularly robust way, but also, less robustly, with RTI reaction time. Furthermore, correlations with cognitive performance were now also found for short time-windows, while global dFC speed variations only showed tendential correlations for long time-windows.

Figure 5A-C shows examples of significant correlations (for the choice of reference modular decomposition made in Figure 4B-C). For short window sizes, decreases in the speed of the dFC module #4 correlated with increase of reaction time in the RTI task (Figure 5A, *p* = 0.044, after Bonferroni correction) and decreases in accuracy on the RVP task for the speed of dFC module #2 (Figure 5B, *p* = 0.0014, after Bonferroni correction). For long window sizes, decreases in the speed of dFC module #4 correlated with an increase in RVP accuracy (Figure 5C, *p* = 0.018, after Bonferroni correction). Table 3 reports several other tendential correlations, for both short and long timewindows.

**Table 3.** Spearman correlations between within-subject variations of modular dFC speed and cognitive change after SD.

| | Short window sizes $\Delta V_{dFC}$ | | | | | Long window sizes $\Delta V_{dFC}$ | | | | |
| --- | --- | --- | --- | --- | --- | --- | --- | --- | --- | --- |
|  | #1 | #2 | #3 | #4 | #5 | #1 | #2 | #3 | #4 | #5 |
| **RTI $\Delta RT$** | — | *-0.67*<br>(*) | — | **-0.90**<br>* | — | — | — | *-0.66*<br>(*) | *-0.73*<br>(*) | — |
| **n-back $\Delta RT$** | — | — | — | — | — | — | — | — | — | *-0.76*<br>(*) |
| **n-back $\Delta acc$** | — | — | — | — | — | — | — | — | *0.78*<br>(*) | — |
| **RVP $\Delta RT$** | — | — | — | — | — | *-0.82*<br>(**) | — | — | — | — |
| **RVP $\Delta acc$** | — | **0.93**<br>** | — | *0.71*<br>(*) | — | — | *0.72*<br>(*) | — | **0.89**<br>* | — |

*RT = Reaction Time; acc = accuracy. Spearman correlation values between within-subject variations of modular dFC speeds (dFC modules #1 to #5, see Figure 4) and within-subject variations of cognitive performance scores. We report values for all significant correlations (bold-face and dark shaded table entry) or tendential correlations. The modular decomposition presented in Figure 4B-C and Figure 5A-C is here used as a reference to compute modular dFC speeds. A \*\* denotes that $p' < 0.01$ ( $p'$ , Bonferroni-corrected p-value, corrected on 2 window size ranges, 5 dFC modules and 5 cognitive tests). We also report (in brackets) significance as assessed from uncorrected p-values to highlight tendential correlations (correlation values in italic and light shaded table entry are significant before correction, i.e. (\*), $p < 0.05$ ; (\*\*), $p < 0.01$ ).*

When considering variations of RSN-restricted dFC speed, instead of modular dFC speed variations, we also found some tendential correlation with cognition (summarized by Table S1). Remarkably, dFC speed restricted to the Ventral Attentional RSN showed a tendential correlation with variation of RTI reaction time and accuracy in the RVP tasks. However, none of these correlations between cognition and RSN-restricted dFC speed survived multiple comparison correction, indicating that decomposition of dFC into unsupervised MC-based modules yield a superior description of relations between dFC and fine cognitive performance variations than a decomposition into expert-based a priori subnetworks of interest.

### Correlations between variations of modular dFC speed and of cognitive performance are robust

The modular decomposition of Figure 4B-C was obtained applying a standard community detection algorithm on the MC matrix of Figure 4A. Such algorithms involve stochastic elements (see *Materials and Methods*) and can, as such, produce different solutions when run multiple times. To verify how robust our conclusions on the relation between modular dFC speeds and cognition were with respect to changes of the detailed decomposition into dFC modules, we extracted a sample of 2000 alternative instances of modular decomposition, starting each time from a different initial condition.

Firstly, we found that the 5 dFC modules described in Figure 4B-C are very robustly retrieved across the various modular decomposition instances. As shown in S9A, a large majority of the extracted decompositions give rise to modules which are very highly correlated with the reference ones. Secondly, by computing correlations between the dFC modular speed variations associated with the alternative instances and cognitive performance, we found patterns of correlations that were highly similar to the ones reported in Table 3 for the reference modular decomposition. As shown in S9B, indeed, we found a mean similarity of ~86% between the speed variations vs cognitive performance variations correlation matrix computed for the reference or the alternative modular decompositions.

As reported in Figure 5D, ~80% of the extracted modular decompositions led to the detection of at least one robustly significant correlation between changes of modular dFC speed and cognitive performance (i.e. after Bonferroni correction). Sometimes even 2 or 3 correlations still significant after multiple comparison correction were found (as for the reference modular decomposition of Figure 4B-C, whose associated significant correlations are highlighted by Figures 6A-C). Among the possible correlations, the ones that were most frequently assessed as significant were correlations with performance on RVP accuracy after SD: overall ~57% of all significant correlations, among which a ~27% of correlations between dFC #2 speed on the short window sizes and RVP accuracy (cf. Figure 5E). This correlation between short-window speed variations of the frontoparietal module dFC #2 and RVP accuracy was by far the most frequent significant correlation, followed by a ~16% of correlations with RVP accuracy with dFC module #4 and a ~15% of dFC module 2 on the long window sizes. We also recorded: a ~11% fraction of significant correlations between dFC #4 modular speed on short window sizes and RTI reaction time; and a ~3% between dFC #3 modular speed on long windows and n-back accuracy. All other correlations were weaker.

In conclusion, the reference modular decomposition of Figure 4B-C, used for the analyses of Figure 5A-C, although particularly fortunate (it is rare to display three robustly significant correlations), is very well representative of general tendencies present in the entire studied set of 2000 alternative modular decompositions.

## Discussion

We found that 24 h of SD systematically induces slowing of dFC on the whole-brain level, i.e. of the speed in which the space of possible functional network configurations are explored. This tendency to slow down, analogous to what previously reported for descent from wakefulness to sleep (El-Baba et al., 2019), is robust at the single-subject level but did not correlate with performance on cognitive tasks. However, when looking at dFC on the local level focusing on metaconnnectivity giving rise to dFC modules, correlation with performance on cognitive tasks was found, notably in sustained attention, despite the large intersubject variability of the actual impact of SD on both cognition and dFC.

Because of the small number of subjects in our sample, the significance of certain correlations with cognitive performance –e.g. global dFC speed with n-back performance variation– or comparisons between conditions –e.g., further slowing down due to sleep staging, cf. Fig. S2– did not survive multiple comparison correction. Therefore, we were not able to investigate the potential physiological mechanisms underlying the observed slowing of dFC speed here. A likely hypothesis (that will have to be explored in future studies) is that the observed slowing may be caused by a SD-induced disruption of slow-wave oscillations (Massimini et al., 2004) or changes in synaptic homeostatic regulation occurring during normal sleep (Tononi & Cirelli, 2014; Diering et al., 2017), that can also be tracked by fMRI studies (Song et al., 2019).

Analogously, extensions of our approach to larger and better adapted cohorts will be needed to study potential confounding sources, such as the impact of having administered a placebo drug to our subjects –which may lead to deviations with respect to a standard resting state condition (Wager & Atlas, 2015)– or different sensitivities to scanner noise (Gaab et al., 2008) before and after SD.

### Modular dFC: a compromise between ignoring time or ignoring space

We first introduced the notion of dFC speed in the companion study by Battaglia et al. (2020). In this article, we confirmed several of the findings of this study, notably the fact that dFC can be seen as a complex random walk in the space of FC network realizations with statistical properties between order and disorder, giving rise to an alternation between “knots” and “leaps”.

The main methodological innovation we introduced here with respect to Battaglia et al. (2020), in order to achieve our aim, was the notion of dFC modules – extracted via the MC analysis – and of modular dFC speeds. While global dFC analyses consider fluctuations of resting-state FC as a whole and quantify how this unique brain-wide network is unspecifically morphing from one time observation to the next, modular dFC analyses explicitly endorse the hypothesis that different functional subnetworks may fluctuate according to independent choreographies, consistent with well-established evidence for regional specialization of brain function, within the context of distributed but separated networks. Modular dFC analyses are thus intermediate between conventional static FC analyses (which describe spatial structure of FC networks at high detail but fully ignore their reconfigurations in time) and the previously introduced global dFC analyses (that track the overall temporal variations of FC in detail while fully ignoring their spatial structure).

### Static MC shapes the structure of dynamic dFC flows

From a methodological perspective, the endeavor of identifying independently fluctuating subnetworks is akin to classic Independent Component Analysis (van de Ven et al., 2004) or decomposition methods relying on link-covariance estimation. We stress indeed that MC is not a novel concept, since edge-based functional connectivities have been previously introduced (Bassett et al., 2014, Brovelli et al., 2017; Faskowitz et al., 2019). However Bassett et al. (2014) or Brovelli et al. (2017) syill considered network components as “hard” reference templates (in a mixture of which wholebrain FC networks are decomposed at any time instant). By contrast, in modular dFC analyses, dFC modules themselves are seen as “soft” dynamic entities, continually reconfiguring in time at changing speeds. In Faskowitz et al. (2019) and even more in Betzel et al. (2019), edge-centric functional connectivities (eFCs) were already used as a way to seize the spatiotemporal complexity of regional co-activation dynamics. Here, dFC as a stochastic exploration of possible functional network configurations provides a transparent geometric interpretation of a sophisticate statistical analysis. Distinct dFC modules in the MC matrix can indeed be seen as being associated to distinct walkers in FC space, performing simultaneously independent random walks. Remarkably, these modules were very robust and largely overlapping between our dataset and MC extracted from an independent control dataset (Figure S5C-E; Termenon et al. 2016). Furthermore, these modules were also evident in the eFC matrix of figure S5B, but MC analysis revealed them even more clearly. Indeed, MC analysis applies a smoothing in estimating the dFC temporal network, by sliding a short averaging window. On the contrary, eFC analysis appears to be noisier, since directly based on instantaneous signal fluctuations. If global dFC speed analyses of Battaglia et al. (2020) describe dFC as a unique flux, according to modular dFC speed analyses, dFC is composed by multiple interlaced streams, smoothly flowing between FC configurations without any rigid separation into discrete FC states (unlike e.g. Allen et al., 2012 or Vidaurre et al., 2016).

It is important to stress here that, given this highly stochastic nature of dFC, sequences of FC matrices do not repeat identically from one subject to the other or even across multiple resting-state sessions for the same subject. In other words, every dFC matrix (Figure 2B top) or, equivalently, its low-dimensional projections via t-SNE embedding (Figure 2B bottom) represents a different random walk snippet realization. However, we expect these dFC realizations to evolve over time respecting a common underlying deterministic structure within a stochastic framework, which is typically characterized by structured flows on manifolds (SFMs; Pillai & Jirsa, 2017) and may be quantified by a common inter-link covariance structure, here described by the MC matrix.

### Modules of dynamic connectivity as cognitive control systems

From a cognitive perspective, separation into dFC modules revealed that the efficiency of cognitive processing in different tasks correlated with the speed of reconfiguration of specific dFC modules but not others. Indeed, while global dFC might be a signature of global cognitive function, confirming previous studies (Wang et al., 2016; Battaglia et al., 2020), modular dFC speeds may show a closer relationship to cognitive performance in different cognitive domains.

Hence, while performance on sustained attention after sleep deprivation as assessed by the RVP task did not correlate with changes of dFC at the whole-brain level, we found a relatively strong correlation of RVP accuracy with dFC speed changes of the frontoparietal dFC module #2. This correlation is furthermore particularly robust being the most frequently found to be significant (cf. Figure 5E) over the large ensemble of alternative dFC modular decompositions that we considered. These results are in line with the functional relevance of ventral and dorsal frontoparietal networks in covert visual attention (Corbetta & Shulman, 2002), the function of the superior parietal cortex and supramarginal gyrus also being associated with visual attention (Coull, Frith, Frackowiak, & Grasby, 1996). Other studies using task-related paradigms have also shown that performance on rapid visual processing, accessing sustained attention, is associated with increased BOLD fMRI activation in frontal and parietal regions (Lawrence, Ross, Hoffmann, Garavan, & Stein, 2003). Moreover, decreased frontoparietal activation after sleep deprivation has been associated with lapses in attention (Chee & Tan, 2010; Chee et al., 2008). Our study is thus in line with previous evidence that frontoparietal networks are sensitive to sleep deprivation and that changes are related to attentional deficits (Muto et al., 2016), as well as for their wide inter-subject variability (Doran et al. 2001, Van Dongen et al. 2004). However, the results of the present report go beyond the previous findings as they show that dynamic reconfigurations in functional connectivity of the frontoparietal network also play a determinant role.

Previous studies have demonstrated that functional networks highly overlapping with task-related networks associated with attentional control are transiently recruited during resting state (Fox et al., 2006). Here, we additionally show that variations after sleep deprivation in the fluidity of connectivity within these intrinsic networks is reduced by sleep deprivation in a way correlating with performance variations in a sustained attention task. We thus reconceptualize these networks as veritable dynamical systems, with fluidly fluctuating internal links, rather than as frozen static networks. Remarkably, the few subjects that improved their RVP performance after sleep deprivation also had an acceleration of their modular speed in the relevant modules, showing that individual cognitive vulnerability or resilience to sleep deprivation, can be tracked by modular dFC.

Finally, while less robust, the significant correlation between dFC #4 modular speed variations and RTI reaction time variations (Figure 5A), can be understood in terms of the involvement of frontal and premotor meta-hubs of dFC #4 module (in part shared with dFC #2 module) in control of action (Passingham 1993). The fact that correlations of modular dFC speeds with RTI and n-back performance are weaker than for RVP may speculatively be linked to a more or less centralized control of cognitive computations involved in the different tasks. Indeed, as previously indicated, the control of sustained attention –probed by the RVP task– relies on the operation of networks which are fairly localized (Corbetta & Shulman, 2002). This is in contrast with, e.g., working memory –probed by the n-back task–, which relies on much more widely distributed networks (Fuster, 1997).”

### dFC and FC modules are qualitatively different entities

It is important to highlight here once again the fundamental difference between FC and dFC modules (cf. Figure 3), although they partially overlap at the level of their hubs and meta-hubs (Figure S6) or when some relation with known RSNs exist (Figure S7). The influence of a hub region on its neighboring regions is quite direct since it causes the fluctuations of adjacent regions’ activity to mirror the ones of its own activity (coordinated node dynamics). On the contrary, a meta-hub region will display in general a wide spectrum of activity correlations with its neighboring regions. For instance, a meta-hub could be strongly correlated with some of its neighbors and very weakly with others. However, these correlations, independently from their strength, will be all boosted up or weakened in a synchronized manner along (coordinated link dynamics). We can use a musical metaphor to illustrate this fundamental difference between hubs and meta-hubs. Both of them can be seen as leading players in an ensemble of musicians. Hubs will force the other musicians to play the same melodic line they play, in unison or following a strictly parallel harmony as in a simple canon. Meta-hubs could be better described as a jazz band leader precisely giving the groove during written sections and then engaging into an improvisation section, in which they still exert an influence on the other players, but quite loosened, allowing divergent soloist lines to emerge in a more spontaneous and less constrained manner. To a signal of the leader however, the improvisation will stop and all the musicians will go back following the prescribed composition. Thus meta-hubs do not necessarily establish strong correlations with other regions (during a jazz improvisation, the various musical lines – although not random – are relatively free) but rather exert a strong modulatory effect on the time-dependent strengths of correlation within their associated functional networks (as when engaging into an improvised from a composed musical section).

Furthermore, the influence of regional nodes when considered as dFC meta-hubs transcends largely the boundaries of the FC networks into which they can be simultaneously acting as hubs. Indeed as revealed by Figure S4, each dFC module includes covarying links that, originating in large part from the module’s specific meta-hubs, diverge then to “touch” — or be incident to — widespread brain regions. Each module is thus incident to nearly the whole brain and, conversely, every region is incident to nearly every module. When a brain region establishes time-dependent correlations with its neighboring regions (“computing” its instantaneous pattern of FC, we could say), it does it under the competing influence exerted by the meta-hubs of multiple dFC modules, as a marionette put into motion by strings pulled simultaneously by different puppeteers. Every meta-hub (“puppeteer”) sends co-modulated incident functional links (“pull strings”) toward nearly every brain region (“marionette”). Thus, localized meta-hubs have a very distributed influence. In this sense, there is no reason to expect an *a priori* correspondence between FC and dFC modules and the imperfect correspondence between dFC and FC modules in Figure S6 should therefore not appear surprising.

More generally, nodes touched by links in a MC meta-module may be at certain times coordinated (when the co-fluctuating links are in a large strength transient) and thus form a FC*(t)* module in the conventional sense. However, they may be poorly coordinated at other times (when the co-fluctuating links are in a weak strength transient) and thus not forming a FC*(t)* module anymore. The link sets that form different meta-modules will fluctuate toward large or weak strengths at different independent times. Therefore, the static meta-modular organization of MC will result in a dynamic modular organization of FC*(t)*, displaying a transient waxing and waning of changing node-level modules. Indeed, FC modules do not invariantly exist at all times but evolve, absorbing and ejecting nodes or by interchanging them with other modules (Schlesinger et al., 2017). Future studies may explore how the “forces” exerted by multiple competing dFC meta-hub controllers control the dynamic reconfiguration of transient FC modules.

Here, we allow ourselves to summarize in a concise glossary (see Table 4) the main metaphors we have introduced to help conceptualizing novel dFC and MC metrics, in the hope to make even more transparent their difference with standard FC approaches.

**Table 4.** dFC and MC glossary.

|  |  |
| --- | --- |
| **dFC stream** | Sequence of Functional Connectivity networks materializing along time.<br>*Or “How the brain dances to the music of time”* |
| **dFC knots or leaps** | Transient slowing down or acceleration of the FC network reconfiguration, smooth without clear discrete states<br>*Or, respectively, “calm pools or rapids along a flowing stream”* |
| **Meta-Connectivity (MC)** | Correlations between the fluctuations of pairs of dynamic functional links<br>*Or, “guidelines for improvisation (whom to tune to the unwritten melody)”* |
| **dFC modules** | Groups of FC links whose fluctuations are correlated between them, but uncorrelated with the ones of links in other groups<br>*Or, “group of players listening to each other in a big jazz band”* |
| **Meta-hubs** | Brain regional nodes whose incident FC links display correlated fluctuations (i.e. centers of FC star subgraphs with strongly meta-connected edges)<br>*Or, “puppet masters”; “players taking the lead in a jazz band”* |

### Conclusion

To summarize, we here introduced a novel framework – modular dFC speed analysis – which allows identifying distinct sets of meta-hub nodes which jointly coordinate the stochastic fluctuations of FC along time in the resting state. Going beyond detection of global effects, we have used our novel analytic framework to correlate speed variations of specific dFC modules to changes in cognitive performance following SD, which vary between subjects. It is already known that sleep deprivation effects express individual phenotypic diversity, which could be explained by genetic differences (Kuna et al. 2012). Furthermore, some of the system-level mechanisms leading to cognitive alterations after SD may be shared with Alzheimer’s disease (Mander et al., 2016). Future studies may exploit the sensitivity of modular dFC analyses to inter-subject differences to track the development of cognitive deficits along Alzheimer’s diseases longitudinal progression. In line with the hypothesis that a superior fluidity of dFC is a proxy for enhanced information processing capabilities (Braun et al., 2018), modular dFC speed could provide a novel marker of the notion of protective “cognitive reserve” (Stern et al. 2009), whose neuroimaging characterizations have been explored more often in terms of task-related activations (Stern, 2017) or structural aspects (Bartrés-Faz & Arenaza-Urquijo, 2011). However they remain less explored at the level of functional connectivity (Martinez et al., 2018) and, especially, concerning its dynamic features, while, on the contrary, which are likely to more closely reflect ongoing neural computations and their efficiency.

## Abbreviations

*rs*: resting-state
*fMRI*: functional Magnetic Resonance Imaging
*BOLD*: blood oxygen level dependent
*FC*: Functional Connectivity
d*FC*: dynamic Functional Connectivity
*MC*: Meta-Connectivity
*RTI*: simple reaction time
*n-back*: working memory task
*RVP*: rapid visual processing task

## Acknowledgements

DB acknowledges support from the Mission for Interdisciplinarity of the CNRS (Infiniti program 2017-2018, “BrainTime”) and from the EU Innovative Training Network “i-CONN” (H2020 ITN 859937). DL has been funded by the Agency of Research of innovation (ANII, Uruguay, grant POS EXT 2015 1 123495). VJ acknowledges funding by the European Union’s Horizon 2020 Framework Program for Research and Innovation under the Specific Grant Agreement No. 785907 (Human Brain Project SGA2). Experimental results have been obtained within the PHARMACOG consortium funded by the European Community’s Seventh Framework Program for the Innovative Medicine Initiative (Grant Agreement no. 115009).

## Supplementary Tables

**Table S1.** Spearman correlations between within-subject variations of RSN-restricted dFC speed and cognitive change after SD.

| | Short window sizes $\Delta V_{dFC}$ | | | | | Long window sizes $\Delta V_{dFC}$ | | | | |
| --- | --- | --- | --- | --- | --- | --- | --- | --- | --- | --- |
| | RTI $\Delta$<br>RT | n-back<br>$\Delta$ RT | n-back<br>$\Delta$ acc | RVP<br>$\Delta$ RT | RVP<br>$\Delta$ acc | RTI $\Delta$<br>RT | n-back<br>$\Delta$ RT | n-back<br>$\Delta$ acc | RVP<br>$\Delta$ RT | RVP<br>$\Delta$ acc |
| RSN Visual | — | — | 0.78<br>(*) | — | — | — | — | — | — | 0.80<br>(**) |
| RSN Somato-motor | — | — | — | — | — | — | — | — | — | — |
| RSN Attention | -0.73<br>(*) | — | — | — | — | -0.84<br>(**) | — | — | — | 0.85<br>(**) |
| RSN Fronto-temporal | — | — | — | — | — | — | — | — | — | — |
| RSN Fronto-<br>parietal | — | — | — | — | — | — | — | — | — | — |
| RSN Default<br>Mode | -0.72<br>(*) | — | — | — | — | -0.84<br>(**) | — | — | — | 0.66<br>(*) |

RT = Reaction Time; acc = accuracy. Spearman correlation values between within-subject variations of RSN-restricted dFC speeds and within-subject variations of cognitive performance scores. We report values for all tendential correlations. A \*\* denotes that $p < 0.01$ ; \*, $p < 0.05$ ( $p$ , Bonferroni-uncorrected $p$ -value). No correlation remained significant after multiple comparison correction.

## Supplementary Figures

**Figure S1.**
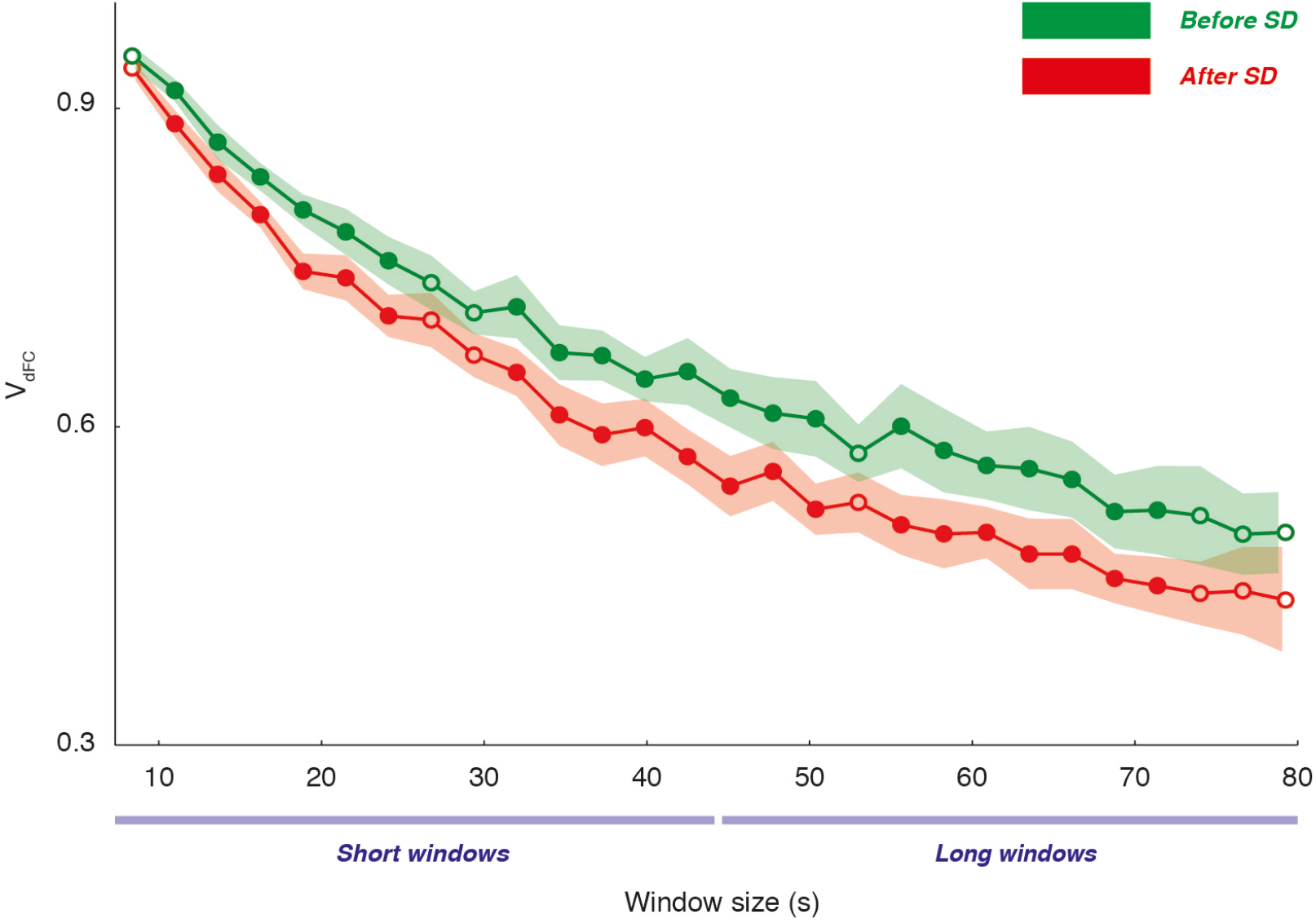
Single window dFC speed analyses. To reduce the number of multiple comparisons and to cope against potential aliasing artifacts, we pool window sizes in two (short and long windows) ranges to estimate typical dFC speed. Nevertheless, the slowing of dFC speed after 24h of SD (red, after SD; green, before SD) is visible even when using individual window sizes. The shaded ranges denote 95% confidence intervals on mean single-window dFC speed. Full (empty) dots denote window sizes for which (prior to Bonferroni correction) the difference between before and after SD would be (not) significant (direct comparison of mean 95% c.i.s).

**Figure S2.**
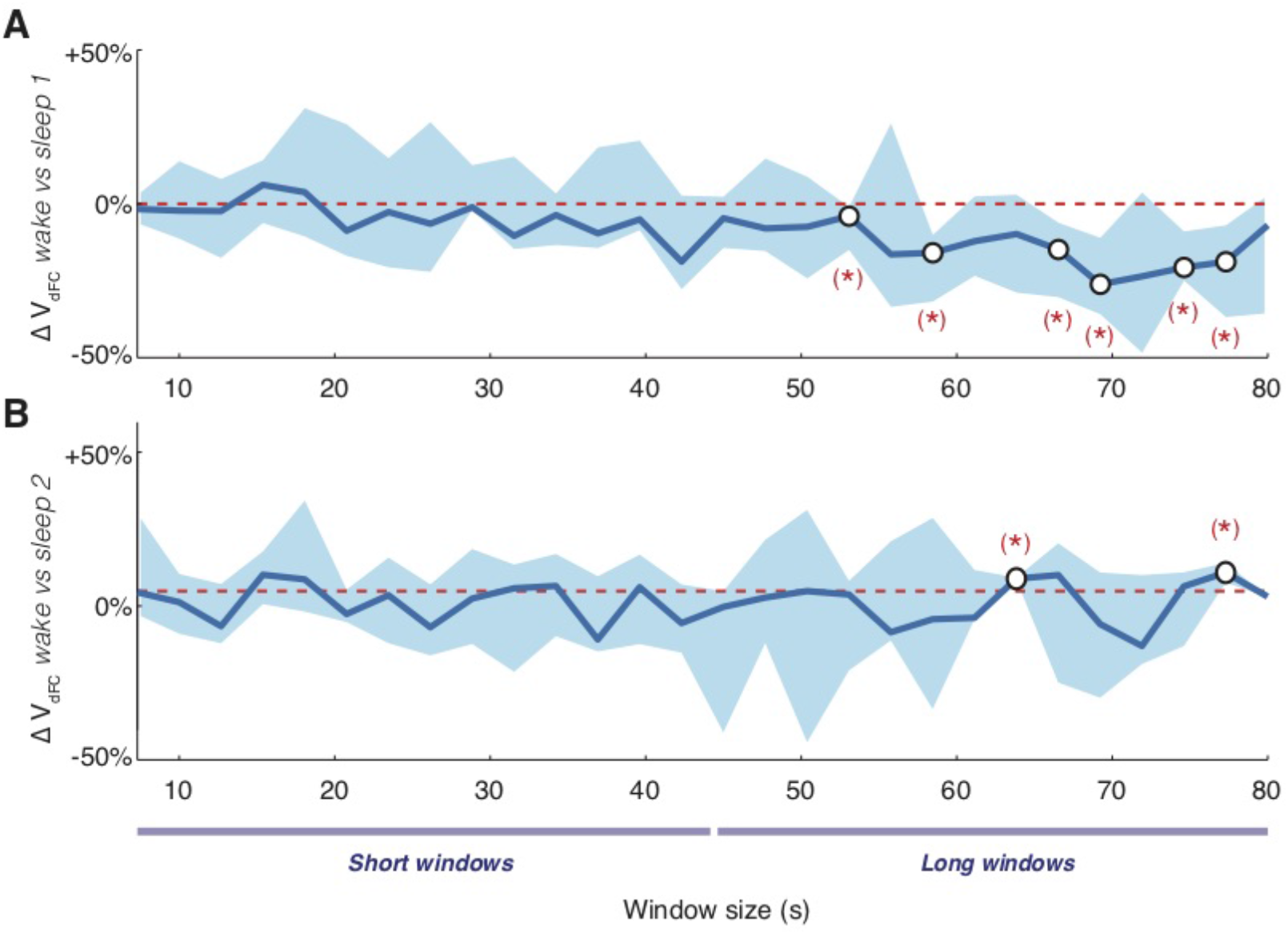
Dependency of global dFC speed on sleep stage. The subjects were monitored for sleep episodes occurring inside the scanner during the resting stare fMRI sessions. Especially after SD, some of the subjects experienced short episodes of sleep. While in the main text Figure 2, we extracted dFC speeds pooling together drowsy with awake epochs, we study here variations in global dFC speed across wake and sleep at increasingly deeper stages (stage N1; stage N2). Within-subject percent variations of global dFC speed across wake and sleep stage 1 are shown in panel **A**, while variations across wake and sleep stage 2 are shown in panel **B**. Given the small duration of sleep episodes for most subjects, no significant differences could be detected for any of the probed window sizes. Only some tendential variations could be detected –speed decrease from wake to sleep stage 1 and speed increase from wake to sleep stage 2– for a few window sizes in the long window range. The symbol (*) denotes *p* < 0.05 (before but not after Bonferroni correction).

**Figure S3.**
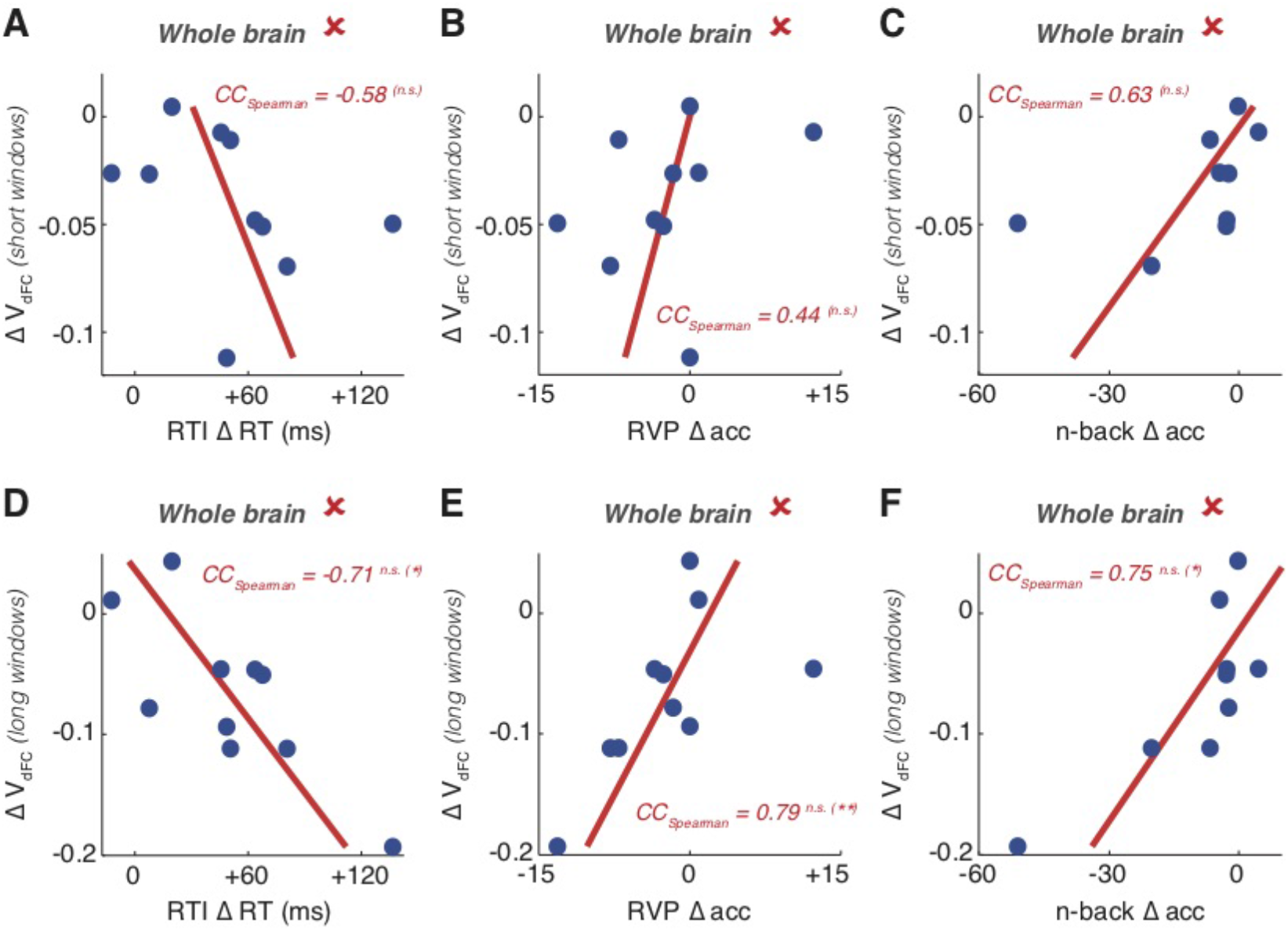
Global dFC speed variations after SD do not correlate with variations of cognitive performance in specific tasks. Scatter plots of the subject-specific variations of (**A** and **D**) RTI reaction time, (**B** and **E**) RVP accuracy and (**C** and **F**) n-back accuracy against the corresponding subject-specific variations of global dFC speed *V_dFC_*, on both short (**A-C**) and long (**D-F**) window sizes. Both cognitive performance and global dFC speed variations are diverse across subjects. However, their correlation is not significant for short window sizes and is only tendential for long window sizes. Stars denote significancy of effect: n.s., not significant; *, *p* < 0.05; **, *p* < 0.01; Bonferroni-corrected values outside brackets and Bonferroni-uncorrected within brackets (tendential correlations).

**Figure S4.**
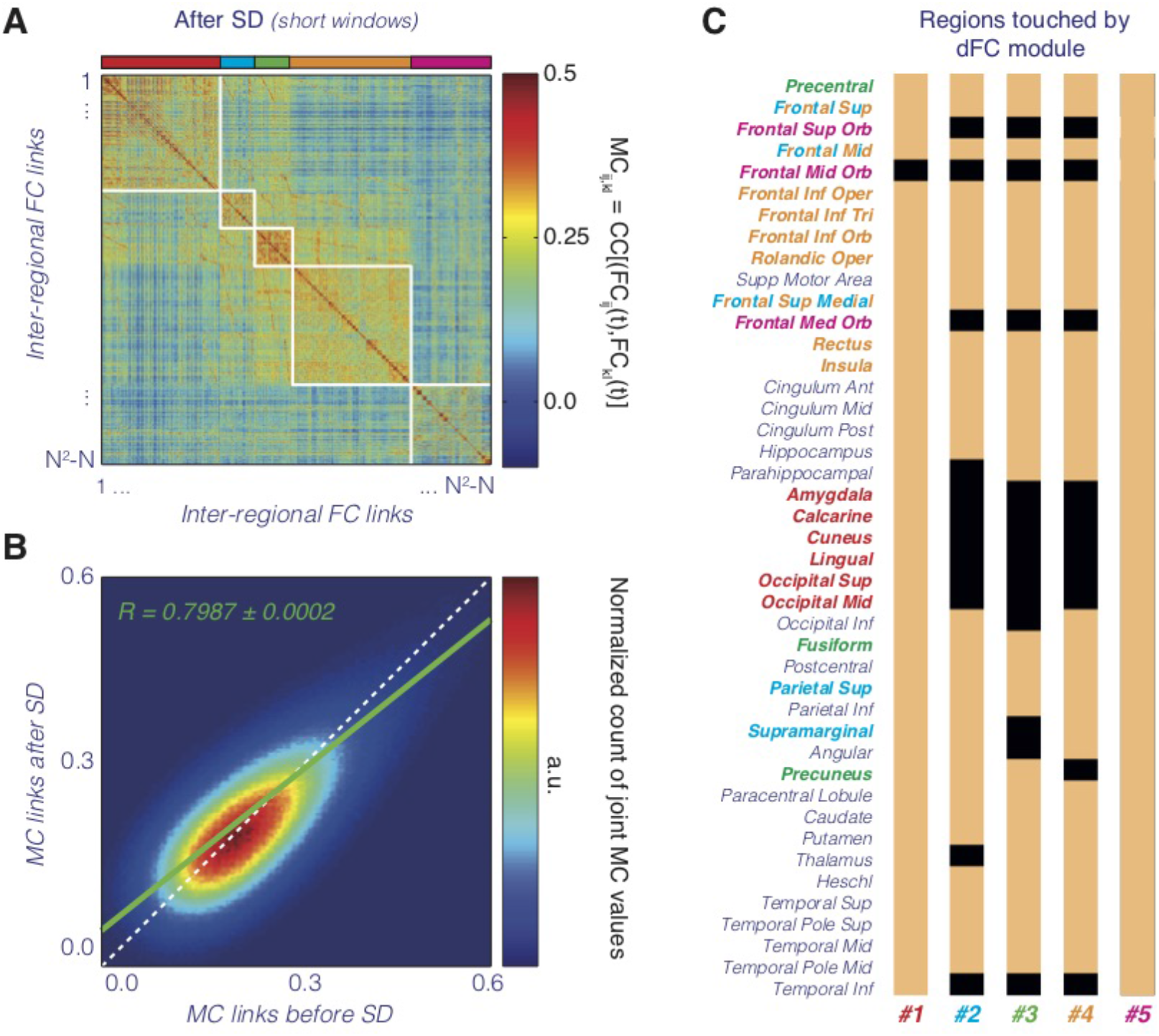
Additional information on MC analysis. **A**. Group-averaged MC matrix for resting state fMRI sessions after SD (cf. Figure 4A for MC before SD). The order of the inter-regional FC links have been rearranged according to modular structure based on MC before SD. No evident variations in modular structure can be detected. **B**. Comparing meta-link strengths before and after SD (entry-wise comparison of MC matrices of Figure 4A and Figure S4A), we find a slight tendency to a decrease in meta-connectivity after SD, as revealed by a slope of fit smaller than 1. **C**. Even if dFC modules have clearly localized meta-hubs (cf. Figure 4B-C), this does not mean that they correspond to clearly localized networks involving just these meta-hubs. We here report for every dFC module all the regions which are incident to at least one FC link belonging to the correspondent dFC meta-module (reminder: meta-modules are communities of links!). A majority of regions are incident to most dFC modules. Therefore dFC modules should not be seen as segregated FC networks but as co-existent systems of influence who independently add up their modulation on the fluctuations of brain-wide FC.

**Figure S5.**
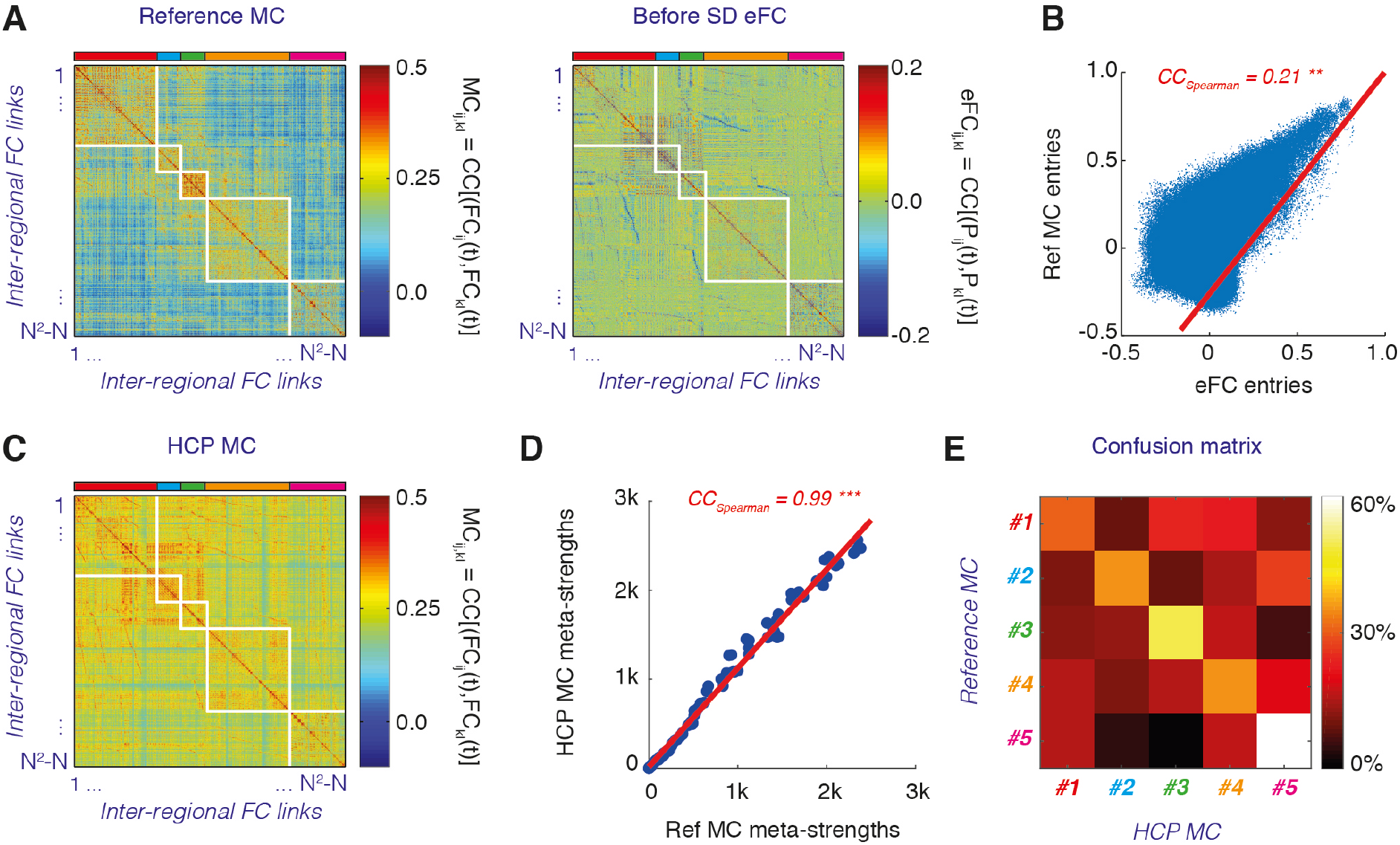
Comparison between reference MC, eFC and control dataset MC. **A**. Shown here (right) is the edge-based Functional Connectivity (eFC), computed according to the definition by Faskowitz et al. (2019) and averaged over all sessions prior to SD. **B.** Scatter plot of the eFC vs the corresponding MC matrix entries. **C.** MC matrix evaluated over a control dataset mediated from the Human Connectome Project (HCP; see Termenon et al., 2016). **D.** Scatter plot of the total MC meta-strengths of different regions in the reference MC vs the HCP-derived MC. The correlation is nearly perfect (bootstrap 95% c.i., 0.993 < CC < 0.996**). E.** Confusion matrix indicating the average overlap between link partitions extracted from the HCP-derived MC matrix of panel (C) with the reference link partition.

**Figure S6.**
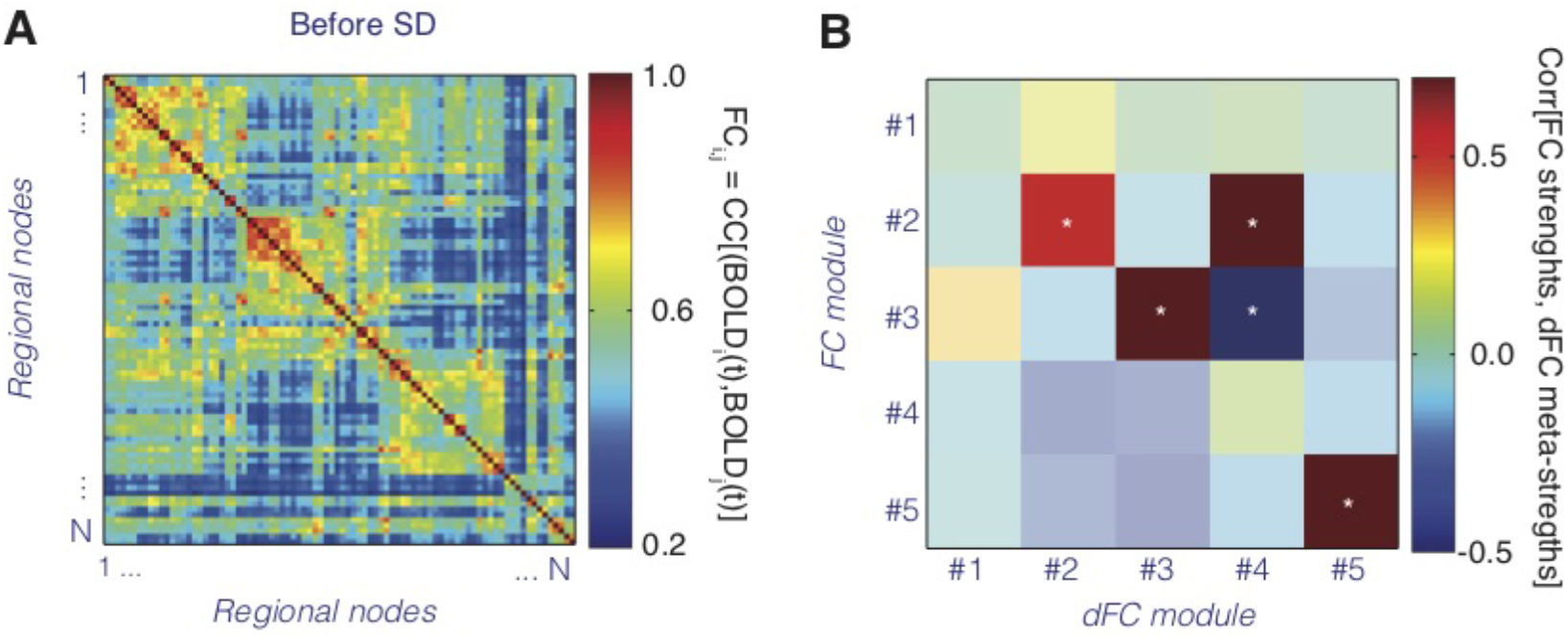
The correspondence between FC and dFC modules is loose. **A**. Group-averaged FC matrix for resting state fMRI sessions before SD. The order of regional nodes have been rearranged to emphasize the modular nature of this time-averaged rs FC. **B.** We extracted 5 modules from the group-averaged FC matrix of panel **A** and compared the participation of the different brain regions into these FC modules with the one that they have into the dFC modules. The matrix shows the correlations between the regional overall FC strengths (restricted to each of 5 FC modules) with the regional overall dFC meta-strengths (restricted to each of the 5 dFC modules). In some cases, there is a good correspondence between FC strengths and dFC meta-strength: e.g. FC and dFC modules #3 and #5 are in good correspondence. However, FC module #2 is related to both dFC modules #2 and #4, failing then to separate them. Furthermore, dFC module #1 is not related to any of the FC modules.

**S7.**
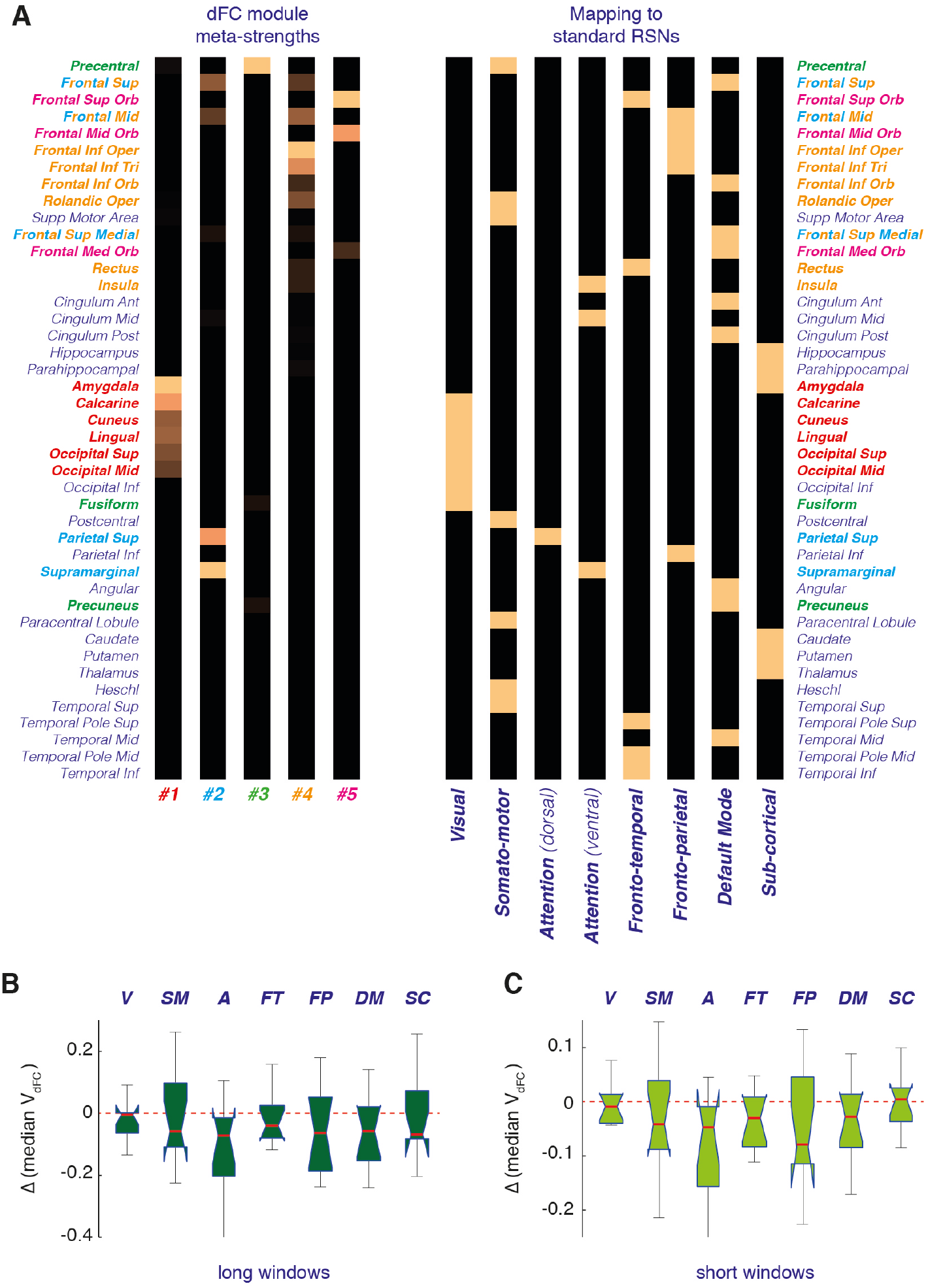
Mapping to Yeo7 Resting State Networks (RSNs). **A**. Reported here to the right is the assignment of the 86 regions of the used AAL atlas to RSNs of the Yeo7 atlas (Yeo et al., 2011), adapted from Amico et al. (2017). Comparison with the integrated meta-strength of different regions within the 5 unsupervised dFC modules extracted from MC analysis reveal an only loose correspondence between dFC module meta-hub systems and RSNs. We combine in the following the unique link included on the Dorsal Attentional Network with the Ventral Attentional network, to give rise to a combined Attentional Network. **B-C**. Within subject variations of RSN-restricted dFC speeds after 24h of SD, for long (panel **B**) and short (panel **C**) window sizes. All modular speeds tend to slow down after SD, however none of these reductions remained significant after applying multiple comparison correction.

**S8.**
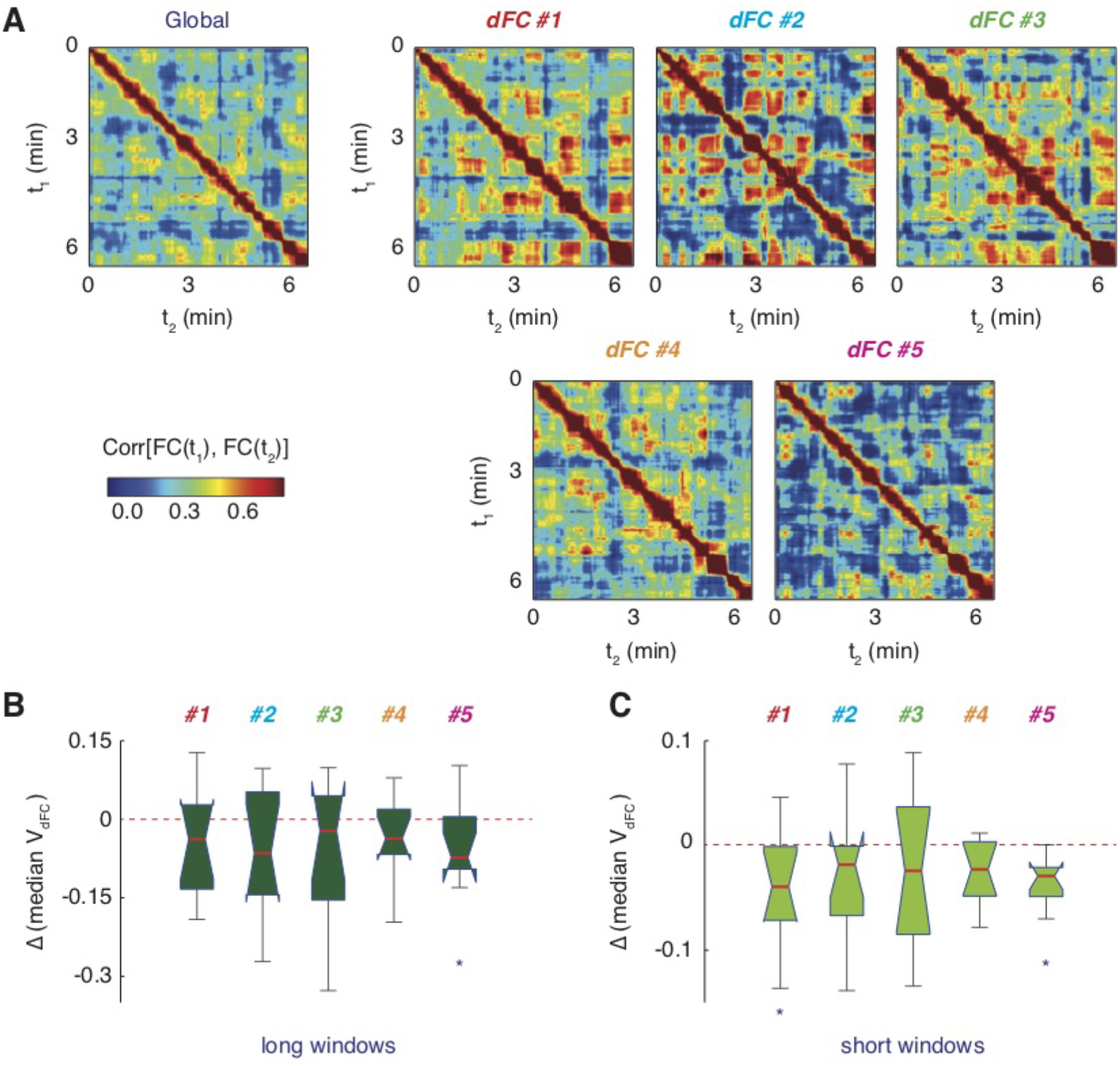
Additional information on modular dFC speed analyses. **A**. The analysis of dFC speed can be restricted to individual modules. We here show representative dFC matrices for each of the 5 reference dFC modules of Figure 4, together with the global dFC matrix (copied from Figure 2B) for comparison (window size of 40 s). The “knots” and “leaps” of these modular dFC matrices are not completely similar, indicating a certain degree of independence between the fluctuations of the distinct dFC modules. **B-C**. Within subject variations of modular dFC speeds after 24h of SD, for long (panel **B**) and short (panel **C**) window sizes. All modular speeds tend to slow down after SD, however significantly only in the case of dFC module #5 for both short and long window sizes and for dFC module #1 for short window sizes. Stars denote significance of effect: *, *p* < 0.05, Bonferroni-corrected values.

**S9.**
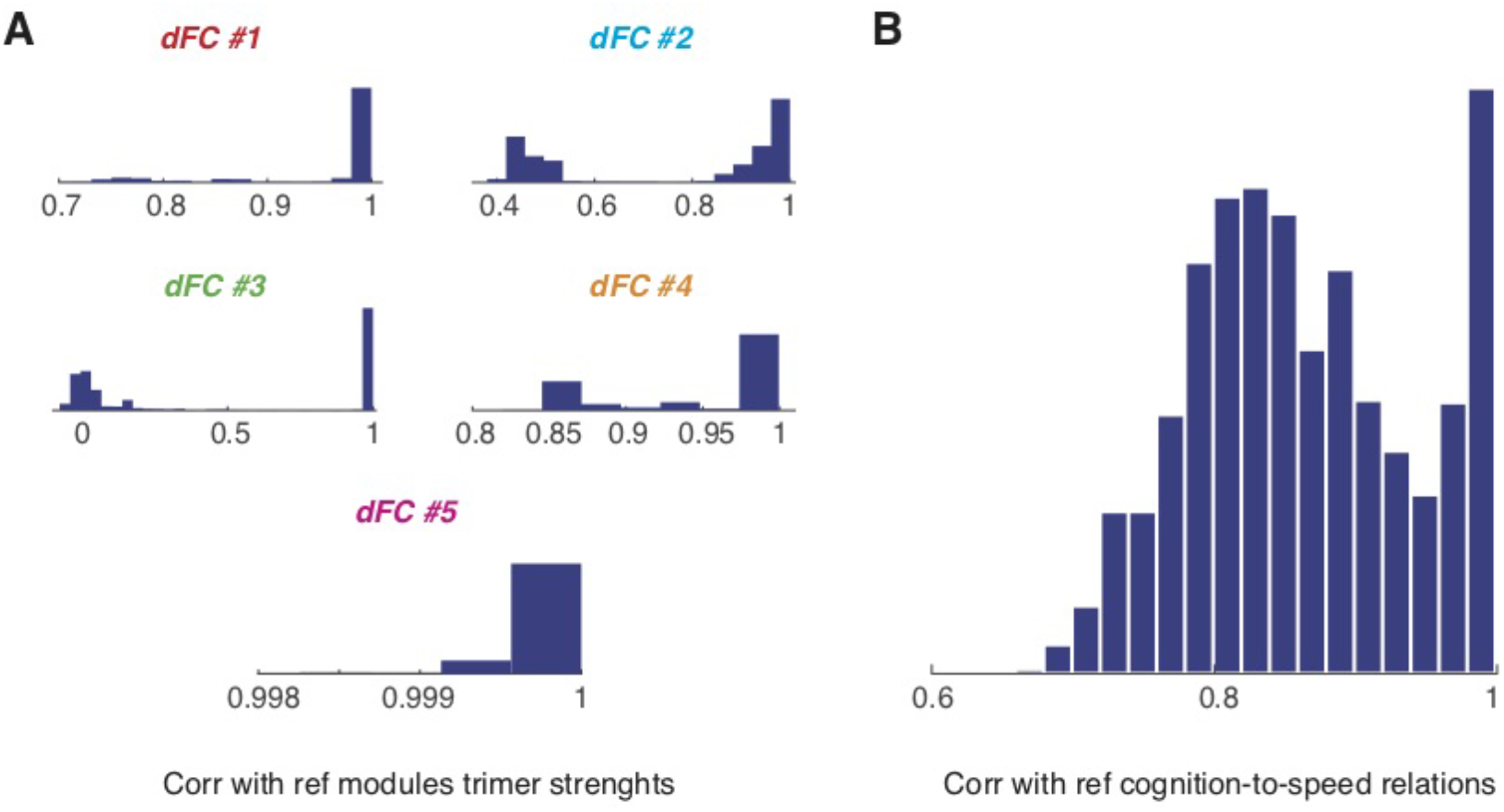
Robustness of dFC modular structure and its correlations with cognition. The algorithm for extracting dFC modules out of the MC matrix of Figure 4A has a stochastic component and produces therefore a different solution for every different run. We chose one specific solution as reference (the one whose modules are portrayed in Figures 5B-C) but studied how different 2000 other instances of modular decomposition from the same MC matrix can be from the chosen reference one. **A**. Histograms of correlation between regional dFC meta-strengths in alternative modular decompositions with the meta-strengths in the chosen reference decomposition. All these histograms have a peak centered on correlation values close to one, indicating that dFC modules highly similar to the chosen reference one are common. In a minority of the solutions, dFC module #3 reduces to a “junk” module without meta-hub regions (hence the secondary peak at near-zero correlation for dFC #3 module alternative-to-reference correlations). When a modular structure includes this degenerate dFC #3 module, then most of the links of the reference dFC #3 are then absorbed by the module analogous to dFC #2 module (hence the secondary peak at near ~0.5 correlation for dFC #2 module alternative-to-reference correlations). Module dFC #5 is nearly perfectly preserved across all modular decomposition instances. **B**. For each of the 2000 alternative modular decompositions we then computed modular dFC speeds and computed correlations between their variations and the variation of cognitive performance. We then computed the similarity between these alternative correlation matrices of modular speed variations vs cognitive performance variations and the one computed for the reference modular decomposition. The histogram of correlations between the alternative correlation matrices and the reference correlation matrix shown in panel **B** strongly peaks toward high similarity values, indicating that correlations between modular dFC speed variations and cognitive performance variations are very robust.

